# Marked Mutants Allow for Rapid Non-Invasive Genotyping of Zebrafish by Visual Inspection Alone

**DOI:** 10.64898/2026.10.29.762891

**Authors:** Sandra Constantinou Juhasz, Toby Shaw-McGrath, Marcus Keatinge

**Affiliations:** Institute for Neuroscience and Cardiovascular Research, University of Edinburgh, UK; UK DRI Edinburgh, University of Edinburgh, UK

## Abstract

Genetic analysis in zebrafish relies heavily on invasive, labour-intensive molecular genotyping. This limits throughput and phenotypic analysis, particularly for homozygous mutants that lack readily detectable phenotypes at early developmental stages. To overcome these limitations, we adapted CRISPR-mediated knock-in strategies to insert complete secondary marker cassettes (including promoter, fluorophore and polyA), directly into coding exons of target loci. This simultaneously disrupts target gene function and permanently labels mutant alleles, which we term Marked Mutants. Crossing heterozygous Marked Mutant alleles targeting the same gene but carrying distinct fluorescent secondary markers linked to each allele, enables unambiguous non-invasive visual identification of wild-type, heterozygous and biallelic mutant progeny as early as 24 hours post fertilisation. To test the scalability and robustness of our approach, we generated eight independent Marked Mutant alleles targeting *mitfa, tdp43* and *tdpl,* using secondary markers comprising hatching gland *(he1.1)*, pancreas *(ins)* or pineal gland *(exorh)* promoters driving expression of GFP, RFP or BFP. We found germline transmission efficiencies comparable to conventional CRISPR mutagenesis, with approximately 30% of fluorescence-selected F0 adults transmitted on-target insertions to the F1 generation. All Marked Mutant alleles behaved as loss-of-function alleles, demonstrating effective disruption of target gene function. By combining four independent Marked Mutant alleles, each carrying a distinct fluorescent secondary marker, we could prospectively identify *tdp43/tdpl* double compound heterozygous embryos by visual inspection alone before phenotypic onset. Only embryos carrying all four Marked Mutant alleles developed the characteristic lethal double-knockout phenotype, while siblings carrying fewer alleles remained phenotypically normal. By eliminating routine PCR genotyping and invasive fin clipping, Marked Mutants simplify colony management, facilitate combinatorial genetics, reduce animal use, and provide a practical platform for rapid non-invasive zebrafish reverse genetics.

## Introduction

A major bottleneck in zebrafish genetic analyses is the reliance on labour-intensive genotyping workflows, which typically require invasive fin clipping of adult fish followed by PCR-based analysis to identify WT, heterozygous and homozygous individuals^1,2^. Genotypes are generally not identified until adult stages, requiring excess genetically modified fish of unneeded genotypes to be raised to adulthood. For example, a heterozygous incross produces approximately 25% homozygous mutants and 25% WT sibling controls, requiring the remaining 50% of heterozygous offspring to be raised until genotyping, despite being unsuitable for the intended experiments. While protocols for genotyping live larval zebrafish have been described, these approaches are either technically demanding, inherently invasive, require anaesthesia or are not readily scalable^3–5^. Moreover, many zebrafish genes are duplicated, with functional redundancy between paralogues often necessitating the generation of double mutants to reveal loss-of-function phenotypes^6–9^. However, an incross of double heterozygotes produces only 1/16 (6.25%) double homozygous mutant offspring. Such experiments require genotyping multiple loci and produce numerous genotype combinations, only a small proportion of which are of experimental interest, substantially increasing the number of animals that must be raised and genotyped. Traditional zebrafish genotyping workflows are particularly restrictive for embryonic-lethal alleles, as homozygous mutants cannot be prospectively identified before phenotypic onset at scale^10^. Moreover, some homozygous mutants may survive embryonic development but struggle to compete with their heterozygous or wild-type siblings^11,12^. Identifying these mutants at scale during embryonic stages would allow them to be separated and given additional care to improve their chances of survival to adulthood. These issues present a major barrier to chemical-genetic rescue and prevention screens, where compounds must be administered before the mutant phenotype emerges, making large-scale screening of genetically defined mutant embryos impractical.

In *Drosophila* genetics, genotyping can be rapid and non-invasive as heterozygous and homozygous individuals can often be identified by visual inspection alone through the use of balancer chromosomes^13^. Balancer chromosomes carry visible markers (such as the *Curly* wing phenotype) and suppress recombination, allowing researchers to track the inheritance of mutant alleles across generations^13^. For example, heterozygous mutants carrying a *Curly*-marked balancer chromosome display curly wings, whereas homozygous mutants lacking the balancer display normal wing morphology^13^.

Long established zebrafish transgenesis strategies frequently incorporate secondary fluorescent markers, such as the green heart marker (Tg cmlc2:GFP), which permit rapid visual identification of transgenic animals through fluorophore expression in discrete anatomical structures as early as 24 hours post fertilisation (hpf)^14,15^. However, conventional transgenesis approaches typically rely on random genomic integration and thus are not targeted. More recently, advances in zebrafish genome editing have enabled the efficient targeting of in frame insertions of fluorophores (such as GFP) into the endogenous N or C termini of target genes^16^. Such gene tagging approaches generate transcriptional reporters or protein fusions which aim to preserve native gene function^16–18^.

Strategies to generate zebrafish mutants where embryos can be genotyped by non-invasive visual inspection have been described, but typically require multiple rounds of insertion, or interfere with live imaging as target gene expression is exchanged for that of a fluorophore^19–21^. This limits their utility in downstream assays requiring live imaging and complicates their application to the study of duplicated genes.

To overcome these challenges, we adapted gene-tagging knock-in methodologies to insert complete secondary marker cassettes (consisting of a promoter, fluorophore coding sequence, transcriptional stop sequence, and polyadenylation signal) directly into coding exons of target genes^16^. Our aim was to simultaneously disrupt the target loci to generate a loss-of-function allele (mutating the gene) while physically linking the mutation to a discrete fluorescent marker for visual identification (marking the mutation). We named these “Marked Mutant” alleles for simplicity to differentiate from traditional zebrafish frameshift/nonsense mutations generated by genome editing/ENU mutagenesis. Crossing two independently generated Marked Mutant (MM) alleles of the same gene, each carrying a distinct secondary marker fluorophore, enables rapid visual identification of biallelic loss-of-function mutants by dual fluorescence, as early as 24hpf, using a conventional fluorescence stereomicroscope, eliminating the need for routine PCR-based genotyping. As proof of principle, we generated Marked Mutant alleles across three different genes, with approximately 30% of fluorescence-selected F0 adults transmitting on-target insertions. This enabled founder identification by screening relatively few selected adults, with the additional advantage of non-invasive visual identification of mutant alleles. To demonstrate the potential of the Marked Mutant approach for combinatorial genetics, we combined four independent Marked Mutant alleles, each carrying a distinct secondary marker, to prospectively identify *tdp43/tdpl* double compound heterozygous embryos by visual inspection before phenotypic onset. Only embryos carrying all four Marked Mutant alleles developed the characteristic lethal double-knockout phenotype, highlighting the potential of this approach to greatly simplify complex genetic analyses in vertebrates.

## Results

### Using secondary markers to disrupt target gene function

We sought to develop a strategy for non-invasive identification of zebrafish larvae carrying biallelic loss-of-function mutations at 24hpf, before the onset of overt phenotypes. We reasoned that PCR-based zebrafish knock-in strategies, previously developed for endogenous gene tagging, could be adapted to insert an entire secondary marker cassette into the coding region of a target gene^16^. Insertion of a secondary marker cassette into the coding sequence should, in principle, introduce premature stop codons irrespective of the reading frame. This would disrupt target gene function while simultaneously providing a permanent fluorescent label in a discrete organ, determined by the promoter driving fluorophore expression (figure 1A and figure 2). Crossing two independently generated Marked Mutant alleles of the same gene, each carrying a visually distinguishable secondary marker, either through different fluorophores expressed in the same tissue or the same fluorophore expressed in different tissues, should enable non-invasive identification of biallelic loss-of-function mutants as early as 24hpf using conventional fluorescence stereomicroscopy (figure 1B).

**Figure 1.**
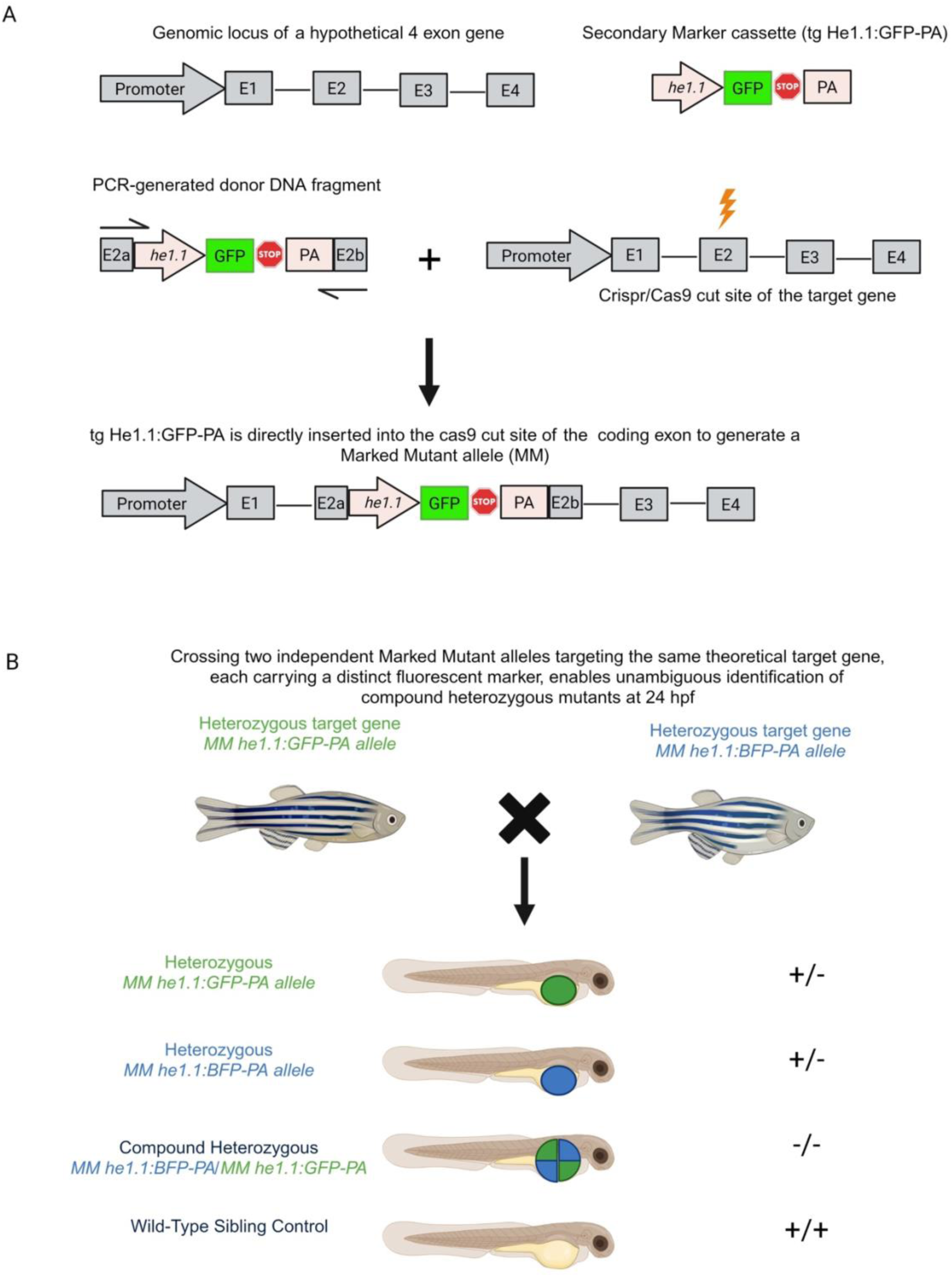
Using secondary markers to simultaneously disrupt and fluorescently label target genes. **(A)** Schematic outlining the Marked Mutant strategy using a hypothetical four-exon target gene. A PCR-generated donor DNA fragment containing a complete hatching gland secondary marker cassette (*he1.1:GFP-PA*) is flanked by 40bp homology arms complementary to sequences either side of the CRISPR/Cas9 cut site in exon 2 of the target gene. The donor DNA is co-injected with a guide RNA targeting the corresponding genomic locus at exon 2. Integration of the secondary marker cassette into the coding exon disrupts target gene function, while the *he1.1* promoter independently drives GFP expression in the hatching gland. **(B)** Crossing two independent Marked Mutant alleles targeting the same gene, each carrying a distinct fluorescent secondary marker (*he1.1:GFP* or *he1.1:BFP*), produces 25% *he1.1*-GFP positive heterozygotes, 25% *he1.1*:BFP positive heterozygotes, 25% compound heterozygotes positive for both *he1.1:GFP* and *he1.1:BFP*, and 25% wild-type siblings. All four genotypes can be distinguished by fluorescence stereomicroscopy as early as 24hpf, enabling rapid, non-invasive identification of biallelic loss-of-function mutants.

**Figure 2.**
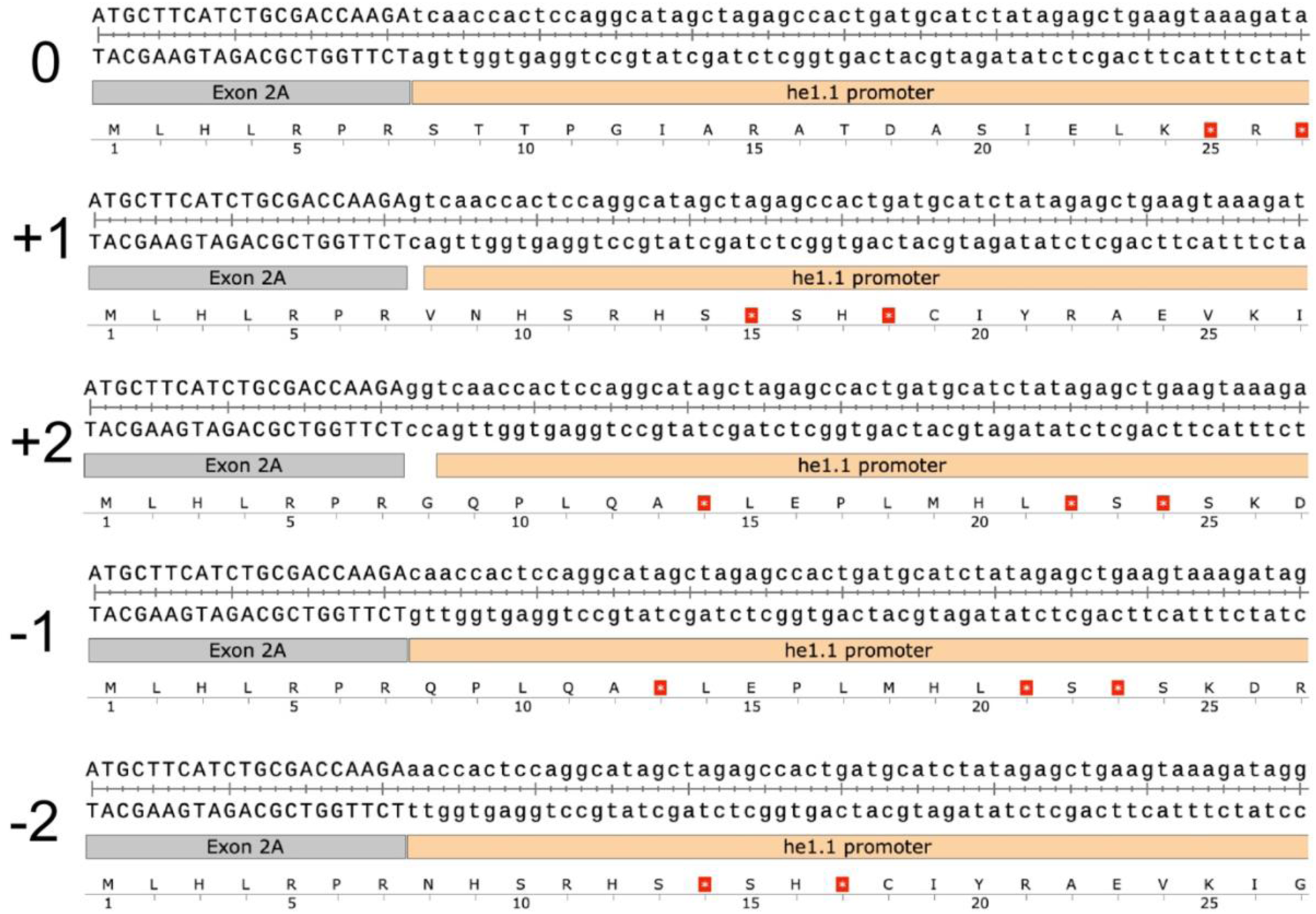
Secondary marker insertion introduces premature stop codons irrespective of reading frame. Schematic illustrating the junction between exon 2 of a hypothetical target gene and the he1.1 promoter of an inserted he1.1 secondary marker cassette. DNA sequences and predicted amino acid translations are shown for five possible insertion junction configurations (0, +1, +2, −1 and −2 bp), representing small insertions or deletions (indels) which could be generated during the integration process. Premature stop codons (red stars) are present within the he1.1 promoter sequence in all five configurations, demonstrating that secondary marker insertion is predicted to disrupt target gene function irrespective of the reading frame and is therefore resistant to small insertion-associated indels.

As a proof of principle, we initially targeted *mitfa*, as homozygous loss of function mutations in this gene lead to the readily identifiable “nacre” phenotype whereby animals appear transparent due to loss of neural-crest-derived melanophores^22^. These attributes made *mitfa* an ideal locus for the initial validation of our Marked Mutant strategy. We co-injected a CrRNA targeting exon 4 of the *mitfa* gene with a PCR product encoding the pineal gland secondary marker (*tg exorh:GFP-PA)*^23^ flanked by 40bp homology arms to each side of the Cas9 cut site (figure 3A). Injections targeted the cell of single cell stage embryos. Approximately 50% of injected F0 embryos exhibited broad GFP fluorescence throughout the pineal gland. This frequency was notably higher than that typically observed using endogenous gene-tagging approaches, in which fluorophores such as GFP are inserted in-frame at the N-or C-terminus of target genes to generate fluorescently tagged endogenous proteins^16^. In our experience, gene tagging approaches generally produce widespread reporter fluorescence in only 1–10% of injected embryos. We reasoned that this increased efficiency reflects a fundamental difference in reporter requirements between the two strategies. Whereas endogenous gene tagging depends on precise in-frame integration of a fluorophore at the target N or C terminal locus to generate a functional fluorescent fusion protein reporter, Marked Mutant secondary marker expression is independent of the target gene reading frame and remains permissive to insertion-associated target gene indels generated during the integration process (figure 2). However, the higher frequency of fluorescence may also reflect episomal expression of the donor DNA, which would not generate functional fluorescent fusion proteins in endogenous gene-tagging approaches^24^. The relative contribution of these mechanisms remains unclear without determining the proportion of fluorescent embryos carrying on-target integrations.

**Figure 3.**
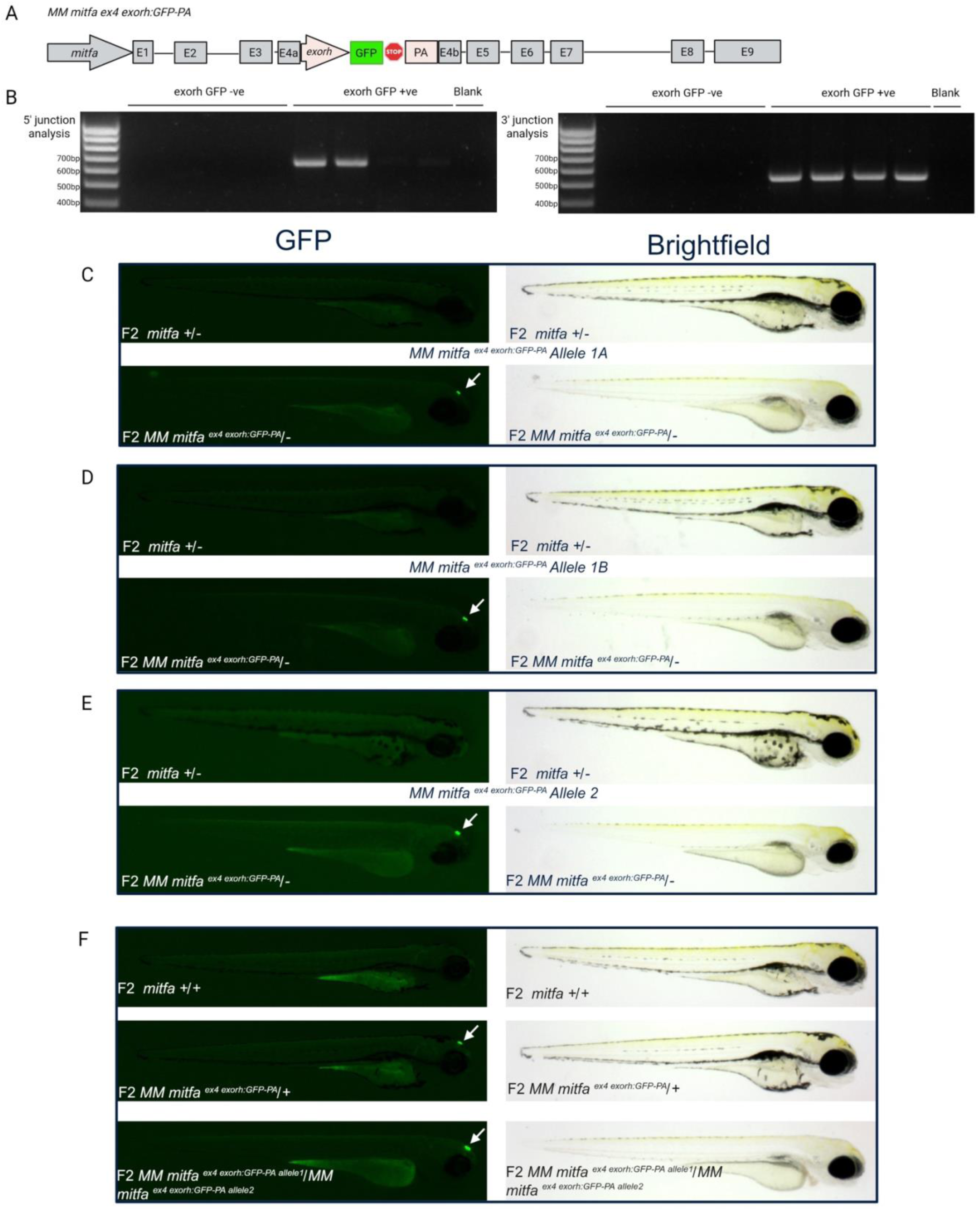
*MM mitfa* alleles are loss of function. **A.** Schematic outlining the pineal gland secondary marker insertion of *exorh:GFP-PA* into exon 4 of the *mitfa* gene to generate the marked mutant allele :*MM mitfa^ex^*^4^ *^exorh:GFP-PA^*. **B.** Junction PCR analysis of F1 from *MM mitfa^ex^*^4^ *^exorh:GFP-PA^* founder 1, denoting at least one intact junction in all *exorh:GFP-PA* positive individuals indicating on target integration. No intact junctions are detected in the GFP negative controls. 50% of GFP positive F1 have both a 5’ and 3’ junction intact whereas 50% only have a 3’ junction. All founder 1 F1 progeny were GFP positive indicating the secondary marker was knocked into the locus of both copies of the endogenous *mitfa* gene, producing two different Marked Mutant alleles which we refer to as allele 1a and allele 1b. **C.** Adult heterozygous F1 *MM mitfa^ex^*^4^ *^exorh:GFP-PA^* allele 1a crossed to conventional *mitfa-/-* produce F2 *MM mitfa^ex^*^4^ *^exorh:GFP^/-* progeny which exhibit the *mitfa-/-* (nacre) phenotype which lack melanophores. This validates the *MM mitfa^ex^*^4^ *^exorh:GFP^* allele 1a as loss of function. **D.** Adult heterozygous F1 *MM mitfa^ex^*^4^ *^exorh:GFP-PA^* allele 1b crossed to conventional *mitfa-/-* produce F2 *MM mitfa^ex^*^4^ *^exorh:GFP^/-* progeny which exhibit the *mitfa-/-* (nacre) phenotype which lack melanophores. This validates the *MM mitfa^ex^*^4^ *^exorh:GFP^* allele 1b as loss of function. **E.** Adult heterozygous F1 *MM mitfa^ex^*^4^ *^exorh:GFP-PA^* allele 2 crossed to conventional *mitfa-/-* produce F2 *MM mitfa^ex^*^4^ *^exorh:GFP^/-* progeny which exhibit the *mitfa-/-* (nacre) phenotype which lack melanophores. This validates the *MM mitfa^ex^*^4^ *^exorh:GFP^* allele 2 as loss of function. **F.** Adult heterozygous F1 *MM mitfa^ex^*^4^ *^exorh:GFP-PA^* allele 1a crossed to adult heterozygous F1 *MM mitfa^ex^*^4^ *^exorh:GFP-PA^* allele 2 ^produce F2 *MM mitfa*^*^ex^*^4^ *^exorh:GFP allele^*^1a^*^/ex^*^4^ *^exorh:GFP allele^*^1a^ *^allele^* ^2^ progeny which exhibit the *mitfa-/-* (nacre) phenotype which lack melanophores. F2 individuals without two copies of a Marked Mutant allele appear WT. This demonstrates that compound heterozygote Marked Mutant alleles can recapitulate homozygous mutant phenotypes.

We raised embryos exhibiting prominent pineal gland GFP fluorescence to adulthood and outcrossed F0 injected animals to wild-type (WT) fish to identify potential founders transmitting the Marked Mutant allele to the F1 generation. Three of four F0 adults produced pineal gland GFP-positive F1 progeny, indicating highly efficient germline transmission of the secondary marker cassette. To confirm targeted integration at the *mitfa* locus, we performed junction PCR analysis on GFP-positive F1 embryos using primer pairs with one primer binding within the secondary marker cassette and the other to flanking endogenous genomic DNA^19^. This generates a PCR product only when the secondary marker is integrated at the intended genomic locus^19^. Progeny derived from all three founders were positive for at least one intact junction, confirming on-target integration at the intended genomic locus (figure 3B). To determine whether these Marked Mutant alleles disrupted gene function, founder fish transmitting the Marked Mutant allele were crossed to *mitfa* -/-loss-of-function mutants. All GFP-positive F1 animals carrying the on target Marked Mutant allele/secondary marker (*MM mitfa^ex^*^4^ *^exorh:GFP^*) displayed the characteristic nacre phenotype when in combination with the conventional *mitfa* mutant allele (*MM mitfa^ex^*^4^ *^exorh:GFP^/-)*, demonstrating that all *MM mitfa^ex^*^4^ *^exorh:GFP^* alleles are loss-of-function.

Notably, founder 1 transmitted the Marked Mutant allele to 100% of its progeny, indicating that secondary marker insertions can be knocked into the germline to homozygosity. Founder 1 produced 50% of progeny with both intact 5’ and 3’ junctions (allele 1a) whilst the other 50% only demonstrated 3’ junctions (allele 1b). The presence of Marked Mutant alleles with only a single intact junction suggested the missing junction was due to either a large insertion or deletion disrupting the amplification of the second junction. To assess whether Marked Mutant alleles remained stable across subsequent generations, we raised F1 *MM mitfa^ex^*^4^ *^exorh:GFP^* heterozygotes carrying one of three alleles: alleles 1a and 1b from founder 1, and an allele from founder 2, which we refer to as allele 2, to adulthood. Each was then crossed again to the conventional *mitfa -/-* mutant line. In all cases, F2 *MM mitfa^ex^*^4^ *^exorh:GFP^/-* progeny exhibited the nacre phenotype, confirming stable inheritance of all Marked Mutant alleles and persistent loss of gene function through the F2 generation (figure 3C-E). To confirm bi-allelic Marked Mutant alleles could reproduce the same phenotypes, we crossed F2 *MM mitfa^ex^*^4^ *^exorh:GFP^* allele 1a heterozygotes with *MM mitfa^ex^*^4^ *^exorh:GFP^* allele 2 heterozygotes. Only *exorh* GFP positive individuals produced the nacre phenotype (figure 3F). All *exorh* GFP negative individuals appeared WT (figure 3F). Genotyping both GFP positive phenotypic, GFP positive WT and GFP negative WT, for the WT *mitfa* locus by PCR confirmed that GFP positive nacre phenotypic individuals lacked a WT copy of the *mitfa* gene. We achieved identical results when crossing F2 *MM mitfa^ex^*^4^ *^exorh:GFP^* allele 1a heterozygotes with F2 *MM mitfa^ex^*^4^ *^exorh:GFP^* allele 1b heterozygotes. This confirms bi-allelic Marked Mutants are loss of function, phenocopying their respective traditional stable mutants. However, as both *mitfa* Marked Mutant alleles carried the same *exorh* secondary marker, heterozygous and biallelic mutants could not be distinguished by fluorescence alone. This highlights the need for distinct secondary markers to enable unambiguous visual identification of biallelic mutants.

### Distinct fluorescent secondary markers enable the generation of complementary *tdp43* Marked Mutant alleles

Many zebrafish genes are duplicated, complicating genetic analysis where paralogues have overlapping functions^25–28^. One such example is *TDP43*, loss of function of which is implicated in amyotrophic lateral sclerosis and frontotemporal dementia (ALS/FTD) in humans^29^. Zebrafish possess two *TDP43* paralogues, *tdp43* and *tdpl*, with robust loss-of-function phenotypes emerging only in double-knockout (DKO) animals, which are lethal by ∼5–6 dpf ^8,9,30^.

Previously, in our hands, the maximum size of exogenous DNA successfully inserted into a target gene using PCR-based gene tagging knock-in approaches was approximately ∼2kb^16^. Importantly, a range of secondary markers of ∼2kb or less, with distinct fluorophores and tissue-specific expression patterns, are available (Table 1)^14,15,23,31,32^. These could be used to generate independent Marked Mutant alleles of the same gene, each carrying a distinguishable secondary marker, allowing heterozygous and biallelic mutants to be identified by fluorescence alone. Furthermore, generating two independent Marked Mutant alleles for each of *tdp43* and *tdpl*, each carrying a distinct secondary marker, would enable the identification of double compound heterozygous embryos through the expression of all four markers. Such individuals should, in theory, phenocopy conventional *tdp43/tdpl* DKOs^9^. We therefore generated complementary Marked Mutant alleles for both genes to test this strategy.

**Table 1.** Secondary markers for rapid, non-invasive visual identification of zebrafish. List of secondary markers that drive fluorophore expression in discrete anatomical structures. Where not previously available, we generated secondary markers expressing red, green or blue fluorescent proteins to increase the range of available marker combinations.

| Gene<br>Promoter | Promoter<br>Length | Expression Pattern |
|---|---|---|
| *he1.1* | ~350bp | Hatching gland from 1-3dpf |
| *cmlc2* | ~500bp | Heart from 24hpf |
| *cryaa* | ~600bp | Lens from 48hpf |
| *exorh* | ~1000bp | Pineal gland from 24hpf |
| *ins* | ~1000bp | Pancreas from 24hpf |

We initially generated marked mutant alleles targeting exon 3 of *tdp43* using the same strategy used to target *mitfa* but replacing the pineal marker with hatching gland-specific secondary marker, driven by the *he1.1* promoter and expressing either GFP or RFP (figure 4A)^14^. This would allow the two mutant alleles to be distinguished by fluorescence, enabling the identification of biallelic mutants by visual inspection alone. The hatching gland represents an advantageous secondary marker for neurological studies because fluorescence becomes detectable early in development (∼20hpf) as a characteristic necklace-like pattern surrounding the yolk sac (figure 4B), then is no longer detectable by ∼4dpf^14^. Consequently, this marker does not interfere with subsequent imaging of the central nervous system (CNS) throughout development.

**Figure 4.**
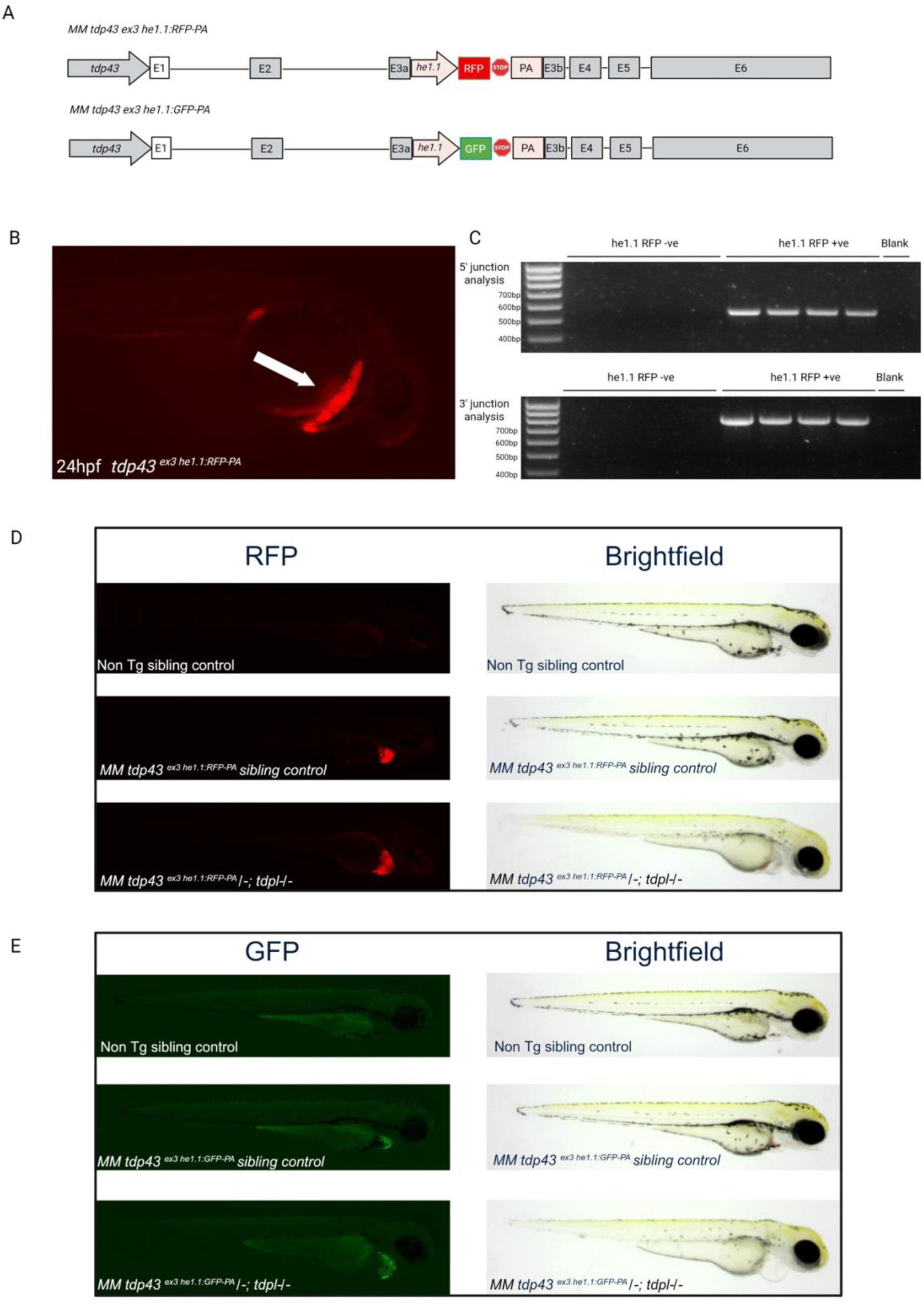
tdp43 Marked Mutations are loss of function. **A.** Schematic outlining the location of *he1.1-RFP-PA* and *he1.1-GFP-PA* secondary markers insertion sites within the *tdp43* coding region. **B.** *MM tdp43^ex^*^3^*^:he^*^1^.^1^*^-RFP-PA^* F1 exhibit RFP fluorescence within the hatching gland (white arrows) at 24hpf allowing for rapid non-invasive visual identification of the Marked Mutant allele. **C.** Junction analysis of *MM tdp43^ex^*^3^*^:he^*^1^.^1^*^-RFP-PA^* founder 1 demonstrates on target integration of the *he1.1-RFP-PA* secondary marker due to the presence of both 5’ and 3’ junction PCR products. Both products are absent from the he1.1-RFP-PA negative sibling controls. **D.** 3dpf F2 Progeny from a F1 *MM tdp43^ex^*^3^*^:he^*^1^.^1^*^-RFP-PA^/+; tdpl +/-* X *tdp43+/-;tdpl +/-* produce F2 *MM tdp43^ex^*^3^*^:he^*^1^.^1^*^-RFP-PA^/-; tdpl -/-* which exhibit the classical DKO phenotype in the form of reduced pigmentation, blood accumulation on the yolk, deformed vasculature and a lack of circulation. The Non Tg sibling control (genotype *tdp43 +/-;tdpl +/-*) and *MM tdp43^ex^*^3^*^:he^*^1^.^1^*^-RFP-PA^* sibling control (genotype *tdp43^ex^*^3^*^:he^*^1^.^1^*^-RFP-PA^/-;tdpl +/-)* demonstrating no phenotype. The DKO phenocopy validates *MM tdp43^ex^*^3^*^:he^*^1^.^1^*^-RFP-PA^* as loss of function. **E.** 3dpf F2 Progeny from a F1 *MM tdp43^ex^*^3^*^:he^*^1^.^1^*^-GFP-PA^/+; tdpl +/-* X *tdp43+/-;tdpl +/-* produce *MM tdp43^ex^*^3^*^:he^*^1^.^1^*^-GFP-PA^/-; tdpl -/-* which exhibit the classical DKO phenotype in the form of reduced pigmentation, blood accumulation on the yolk, deformed vasculature and a lack of circulation. The Non Tg sibling control (genotype *tdp43 +/+;tdpl -/-*) and *MM tdp43^ex^*^3^*^:he^*^1^.^1^*^-GFP-PA^* sibling control (genotype *tdp43^ex^*^3^*^:he^*^1^.^1^*^-GFP-PA^/-;tdpl +/+)* demonstrating no phenotype. The DKO phenocopy validates *MM tdp43^ex^*^3^*^:he^*^1^.^1^*^-GFP-PA^* as loss of function.

Injection of both *he1.1*-RFP-PA and *he1.1*-GFP-PA donor cassettes resulted in widespread hatching gland fluorescence in approximately 50% of injected embryos. Fluorescent F0 animals with prominent hatching gland expression were raised to adulthood and crossed to WT fish to identify founders transmitting Marked Mutant alleles through the germline. For the *MM tdp43^ex^*^3^*^:he^*^1^.^1^*^-RFP-PA^* allele, 3 of 8 F0 adults produced F1 progeny with RFP-positive hatching glands (figure 4B). Junction PCR analysis confirmed that all F1-positive embryos carried on-target integrations for at least one junction from all 3 F0 founders (figure 4C). Because TDP43 deficiency in zebrafish produces robust phenotypes only in the context of combined *tdp43* and *tdpl* DKO, we next assessed whether the *tdp43* Marked Mutant allele behaved as a functional null in this background. Zebrafish *tdp43;tdpl* DKO embryos first develop phenotypes at approximately 2dpf, characterized by reduced pigmentation, impaired or absent blood flow and accumulation of blood within the yolk sac despite the presence of a beating heart^9^. Subsequently, embryos develop severe morphological abnormalities, progressive muscle atrophy and lethality by ∼5-6dpf.

To test the functionality of the Marked Mutant allele, we crossed a *MM tdp43*^ex3:he1.1-RFP-PA^ founder to a conventional *tdpl+/-* loss-of-function line to generate F1 *tdp43 +/MM tdp43^ex^*^3^*^:he^*^1^.^1^*^-RFP-PA^;tdpl+/-* adults. These animals were subsequently crossed to conventional *tdp43+/-;tdpl+/-* double heterozygotes to generate F2 *tdp43^ex^*^3^*^:he^*^1^.^1^*^-RFP-PA^/-;tdpl-/-* embryos. All F2 *tdp43^ex^*^3^*^:he^*^1^.^1^*^-RFP-PA^ /-;tdpl-/-* phenocopied the established *tdp43;tdpl* DKO phenotype, exhibiting reduced pigmentation, severely reduced or absent blood flow, progressive deformity, muscle atrophy and lethality (figure 4D)^9^. These findings confirm that the *MM tdp43^ex^*^3^*^:he^*^1^.^1^*^-RFP-PA^* is a loss-of-function allele.

We next generated an independent *tdp43^ex^*^3^*^:he^*^1^.^1^*^-GFP-PA^* allele using the same strategy. Screening of 10 potential F0 founders identified 3 F0 transmitting GFP-positive hatching gland expression to the F1 generation; however, junction PCR analysis demonstrated that only one founder carried an intact on-target integration. This observation highlights the potential for secondary marker cassettes to integrate at off-target loci and underscores the importance of junction analysis during founder validation. The on target *tdp43^ex^*^3^*^:he^*^1^.^1^*^-GFP-PA^* founder was crossed onto a *tdpl+/-* background as before to generate F1 *MM tdp43^ex^*^3^*^:he^*^1^.^1^*^-GFP-PA^/+*;*tdpl+/-* animals. These were crossed to conventional *tdp43+/-;tdpl+/-* double heterozygotes to generate F2 *tdp43^ex^*^3^*^:he^*^1^.^1^*^-GFP-PA^ /-;tdpl-/-* embryos. All *tdp43^ex^*^3^*^:he^*^1^.^1^*^-GFP-PA^ /-;tdpl-/-* recapitulated the canonical *tdp43;tdpl* DKO phenotype, displaying reduced pigmentation, impaired circulation, accumulation of blood on the yolk sac, muscle atrophy and lethality by ∼5dpf (figure 4E), confirming that *MM tdp43^ex^*^3^*^:he^*^1^.^1^*^-GFP-PA^* is also a loss-of-function allele^9^. In both cases, individuals that retained at least one WT allele of *tdp43* or *tdpl*, or lacked the fluorescent Marked Mutant allele did not exhibit the DKO phenotype. This confirms that both Marked Mutant *tdp43* alleles are loss of function. Importantly, these two independent alleles carry distinct GFP or RFP hatching gland secondary markers, allowing them to be distinguished by fluorescence alone. Generating two additional *tdpl* Marked Mutant alleles with distinguishable secondary markers would therefore enable the identification of *tdp43/tdpl* double compound heterozygous embryos through the combined expression of all four markers.

### Generating complementary *tdpl* Marked Mutant alleles towards four-marker double-knockout identification

Having generated two complementary *tdp43* Marked Mutant alleles, we next sought to generate independent *tdpl* Marked Mutant alleles to enable the identification of DKO using four distinct secondary markers. We also wanted to determine whether target gene function could be disrupted using different secondary markers, cassette orientations and insertion sites. We therefore generated *tdpl* Marked Mutant alleles targeting exon 1 using insulin promoter-driven GFP and BFP secondary markers, including a BFP cassette inserted in the reverse-complement (RC) orientation. Additionally, we generated Marked Mutant alleles using pineal gland (exorh)-driven BFP and GFP secondary markers inserted in the reverse-complement orientation into exons 1 and 2, respectively. We then assessed whether these different cassette designs successfully disrupted *tdpl* function.

We initially targeted exon 1 of *tdpl* using insulin promoter-driven GFP and BFP secondary markers, allowing mutant alleles to be identified by pancreatic fluorescence^31^ (figure 5A). Exon 1 of *tdpl* was targeted using either *ins:GFP-PA* or *ins:BFP-PA* donor cassettes and F0 embryos with widespread GFP or BFP signal in the pancreas were raised for founder screening. Of note, the insulin promoter produced additional fluorescence in the hatching gland and a subset of neurons, which may reflect transient episomal expression of the injected construct. Similar ectopic expression has previously been reported for modified insulin promoter sequences^31^.

**Figure 5.**
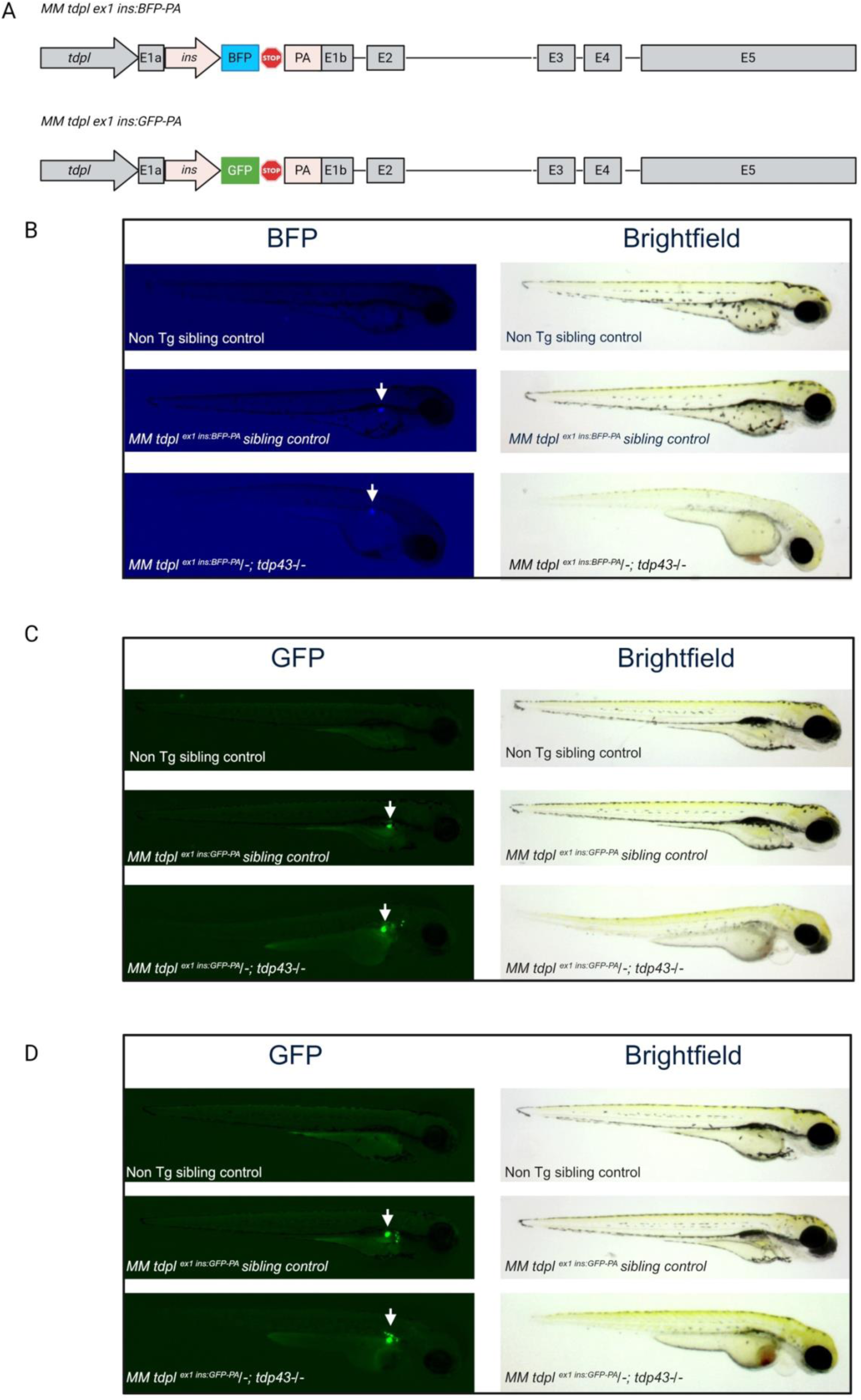
MM tdpl _ex1:ins-BFP-PA_, MM tdpl ^ex1:ins-GFP-PA allele1^ and MM tdpl ^ex1:ins-GFP-PA allele2^ are loss of function. **A.** Schematic outlining the location of *ins-BFP-PA* and *ins-GFP-PA* secondary marker insertion sites within the *tdpl* coding region. **B.** 3dpf F2 progeny from a F1 *tdp43 +/-;MM tdpl ^ex^*^1^*^:ins-BFP-PA^/+* X *tdp43+/-;tdpl +/-* cross produce *MM tdpl ^ex^*^1^*^:ins-BFP-PA^/-; tdp43 -/-* which exhibit the classical DKO phenotype in the form of reduced pigmentation, blood accumulation on the yolk, deformed vasculature and a lack of circulation. The Non Tg sibling control (genotype *tdpl +/+;tdp43 +/-*) and *MM tdpl ^ex^*^1^*^:ins-BFP-PA^* sibling control (*MM tdpl ^ex^*^1^*^:ins-BFP-PA^/+; tdp43 +/-)* demonstrating no phenotype. The DKO phenocopy validates *MM tdpl ^ex^*^1^*^:ins-BFP-PA^* as loss of function. **C** 3dpf F2 progeny from a F1 *tdp43 +/-;MM tdpl ^ex^*^1^*^:ins-GFP-PA allele^*^1^*/+* X *tdp43+/-;tdpl +/-* cross produce *MM tdpl ^ex^*^1^*^:ins-GFP-PA allele^*^1^*/-; tdp43 -/-* which exhibit the classical DKO phenotype in the form of reduced pigmentation, blood accumulation on the yolk, deformed vasculature and a lack of circulation. The Non Tg sibling control (genotype *tdpl +/-;tdp43 +/-*) and *MM tdpl ^ex^*^1^*^:ins-GFP-PA allele^*^1^ sibling control (*MM tdpl ^ex^*^1^*^:ins-GFP-PA allele^*^1^*/-; tdp43 +/-)* demonstrating no phenotype. The DKO phenocopy validates *MM tdpl ^ex^*^1^*^:ins-GFP-PA allele^*^1^ as loss of function. **D.** 3dpf F2 progeny from a F1 *tdp43 +/- ;MM tdpl ^ex^*^1^*^:ins-GFP-PA allele^*^2^*/+* X *tdp43+/-;tdpl +/-* cross produce *MM tdpl ^ex^*^1^*^:ins-GFP-PA allele^*^2^*/-; tdp43 -/-* which exhibit the classical DKO phenotype in the form of reduced pigmentation, blood accumulation on the yolk, deformed vasculature and a lack of circulation. The Non Tg sibling control (genotype *tdpl +/-;tdp43 +/+*) and *MM tdpl ^ex^*^1^*^:ins-GFP-PA allele^*^2^ sibling control (*MM tdpl ^ex^*^1^*^:ins-GFP-PA allele^*^2^*/+; tdp43 -/-)* demonstrating no phenotype. The DKO phenocopy validates *MM tdpl ^ex^*^1^*^:ins-GFP-PA allele^*^2^ as loss of function.

Germline-transmitting founders (with minimal off target florescence) carrying validated on-target integrations were recovered for both alleles, with 3 of 7 F0 adults transmitting *MM tdpl^ex^*^1^*^:ins-GFP-PA^* and 1 of 5 transmitting *MM tdpl^ex^*^1^*^:ins-BFP-PA^* to the F1 generation. Junction PCR analysis confirmed on-target insertion in these founder lines. To determine whether these alleles were loss-of-function, we crossed two *MM tdpl^ex^*^1^*^:ins-GFP-PA^ alleles* and the single *MM tdpl^ex^*^1^*^:ins-BFP-PA^* allele onto a conventional *tdp43+/-* background and raised to adulthood. Subsequent crossing of the F1 with *tdp43+/-;tdpl+/-* generated F2 embryos carrying either *tdp43-/-*; *MM tdpl^ex^*^1^*^:ins-BFP-PA^* /*-* or *tdp43-/-*; *MM tdpl^ex^*^1^*^:ins-GFP-PA^* /- genotypes. In all cases, *tdp43-/-*; *MM tdpl^ex^*^1^*^:ins-BFP-PA^* /*-* and *tdp43-/-*; *MM tdpl^ex^*^1^*^:ins-GFP-PA^* /- phenocopied the established *tdp43;tdpl* DKO phenotype, displaying reduced pigmentation, impaired circulation, progressive deformity and embryonic lethality (figure 5B-D). Individuals that did not carry either the Marked Mutant fluorescent allele, or carried a single WT *tdp43* or *tdpl* allele did not exhibit the phenotype. This confirmed that both marked mutant *tdpl* alleles are loss of function.

We then generated an additional marked mutant *tdpl* allele in which the *ins:BFP-PA* cassette was inserted in the reverse-complement (RC) orientation (Figure 6A). Screening identified 1 of 7 injected F0 adults transmitting an on-target integration of the *MM tdpl ^RC ex^*^1^*^:ins-BFP-PA^* allele to the F1 generation. Phenotypic analysis confirmed that this allele behaved as a loss-of-function mutant (figure 6B). We then targeted a pineal gland reporter cassette to *tdpl* exon 1 and the more distal exon 2 (figure 7A) to test if different secondary markers and insertion sites could cause loss of *tdpl* function. Germline-transmitting founders carrying validated on-target insertions were recovered for both *MM tdpl ^RC ex^*^1^*^:exorh-BFP-PA^* and *MM tdpl ^RC ex^*^2^*^:exorh-GFP-PA^* alleles, with 3 of 11 and 1 of 4 injected F0 adults producing positive F1 progeny, respectively. Both alleles were confirmed as loss-of-function mutants (figure 7B-C).

**Figure 6.**
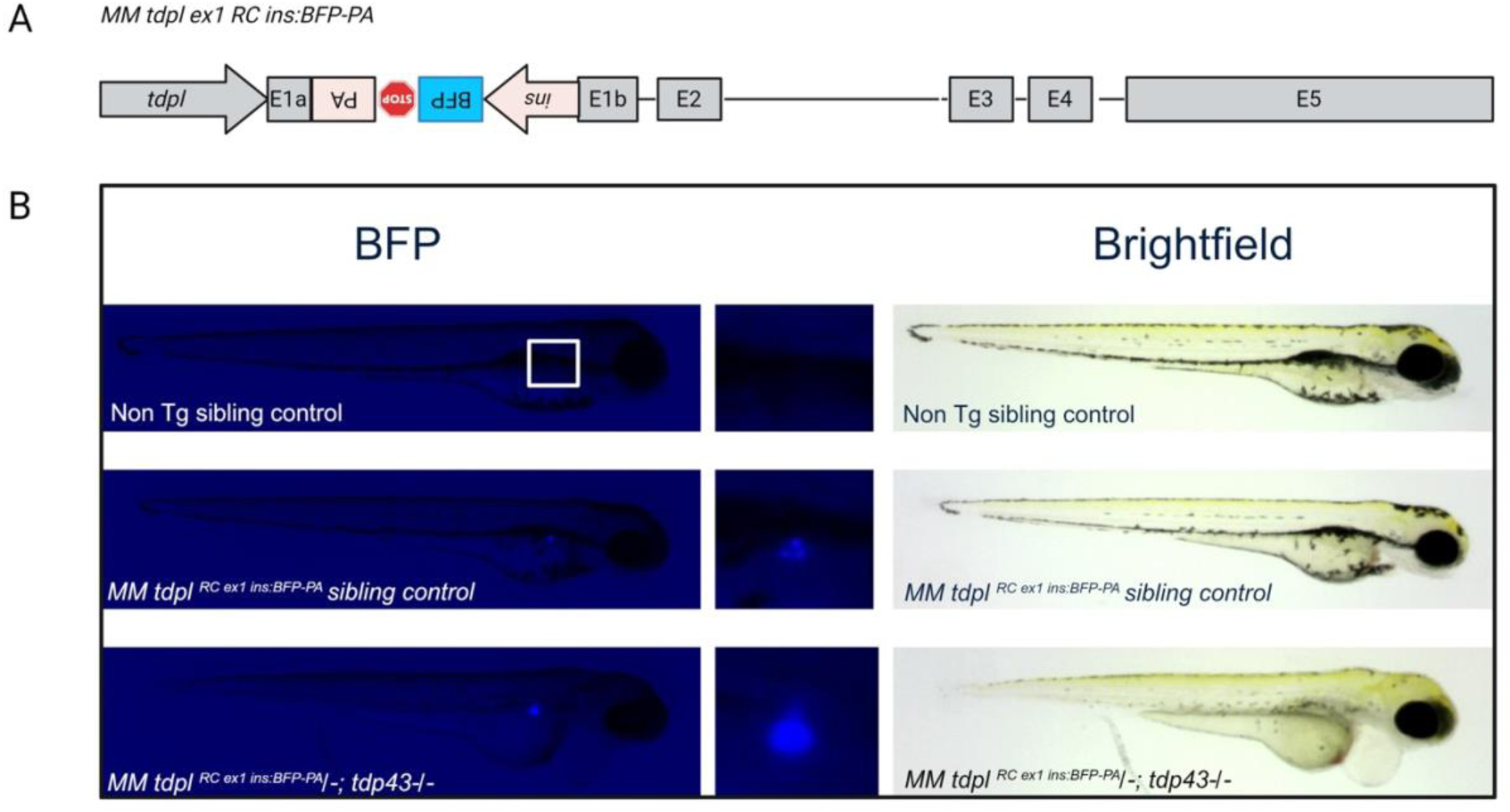
MM tdpl ^RC ex1:ins-BFP-PA^ is loss of function. **A.** Schematic outlining the location of *RC ins-BFP-PA* secondary marker insertion site within the *tdpl* coding region. **B.** 3dpf F2 progeny from a F1 *tdp43 +/- ; MM tdpl ^RC ex^*^1^*^:ins-BFP-PA^ /+* X *tdp43+/-;tdpl +/-* cross produce *MM tdpl ^RC ex^*^1^*^:ins-BFP-PA^ /-; tdp43 -/-* which exhibit the classical DKO phenotype in the form of reduced pigmentation, blood accumulation on the yolk, deformed vasculature and a lack of circulation. The Non Tg sibling control (genotype *tdpl +/-;tdp43 +/+*) and *MM tdpl ^RC ex^*^1^*^:ins-BFP-PA^* sibling control (*MM tdpl ^RC ex^*^1^*^:ins-BFP-PA^ /-; tdp43 +/+)* demonstrating no phenotype. The DKO phenocopy validates *MM tdpl ^RC ex^*^1^*^:ins-BFP-PA^* as loss of function.

**Figure 7.**
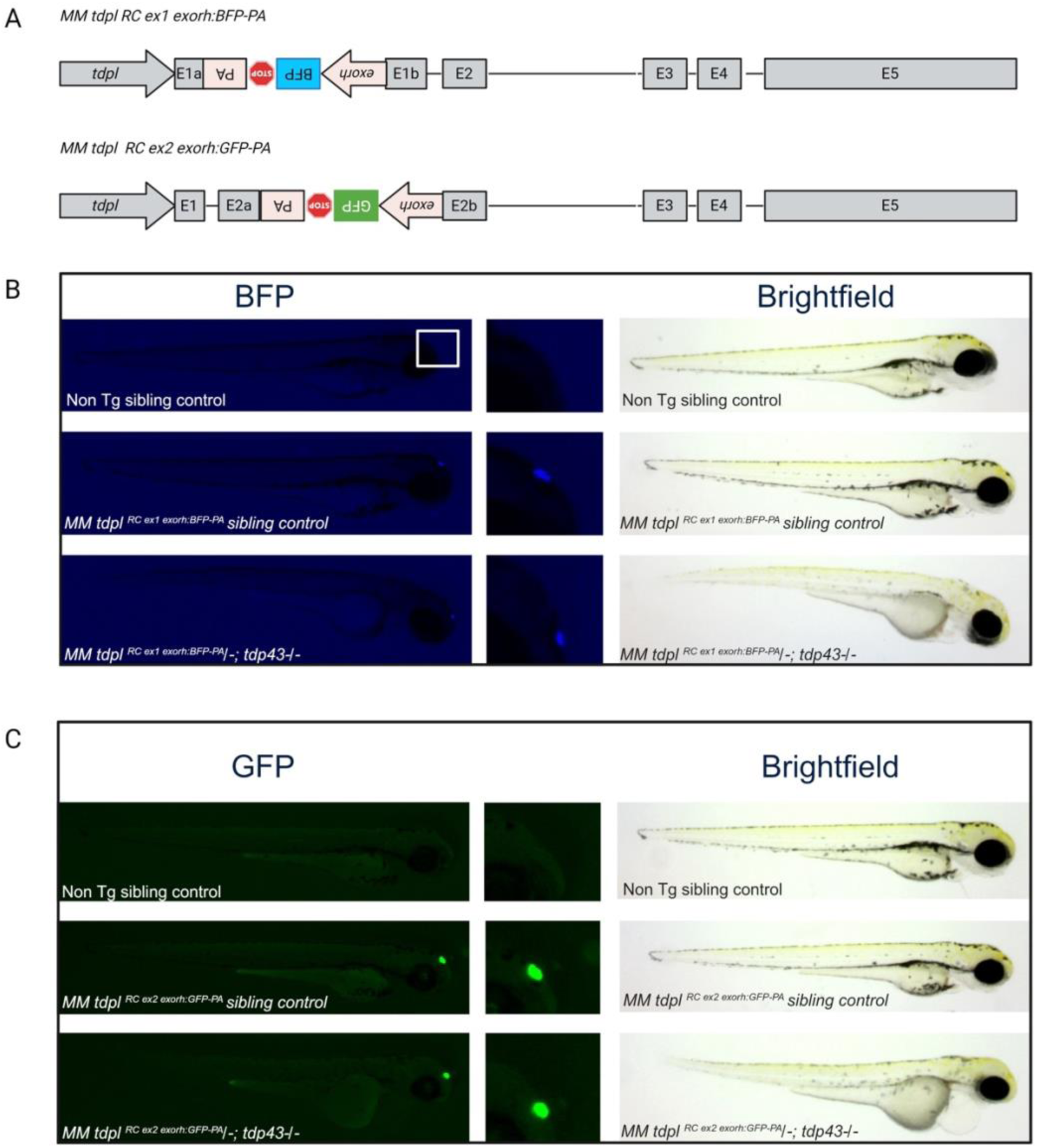
MM tdpl ^RC ex1:exorh-BFP-PA^, MM tdpl ^RC ex2:exorh-GFP-PA^ are loss of function. **A.** Schematic outlining the location of *RC exorh-BFP-PA* and *RC exorh-GFP-PA* secondary marker insertion sites within the *tdpl* coding region. **B.** 3dpf F2 progeny from a F1 *tdp43 +/- ; tdpl ^RC ex^*^1^*^:exorh-BFP-PA^/+* X *tdp43+/-;tdpl +/-* cross produce *MM tdpl ^RC ex^*^1^*^:exorh-BFP-PA^/-; tdp43 -/-* which exhibit the classical DKO phenotype in the form of reduced pigmentation, blood accumulation on the yolk, deformed vasculature and a lack of circulation. The Non Tg sibling control (genotype *tdpl +/+;tdp43 +/-*) and *MM tdpl ^RC ex^*^1^*^:exorh-BFP-PA^* sibling control (*MM tdpl ^RC ex^*^1^*^:exorh-BFP-PA^ /-; tdp43 +/+)* demonstrating no phenotype. The DKO phenocopy validates *MM tdpl ^RC ex^*^1^*^:exorh-BFP-PA^* as loss of function. **C** 3dpf F2 progeny from a F1 *tdp43 +/- ; MM tdpl ^RC ex^*^2^*^:exorh-GFP-PA^/+* X *tdp43+/-;tdpl +/-* cross produce *MM tdpl ^RC ex^*^2^*^:exorh-GFP-PA^ /-; tdp43 -/-* which exhibit the classical DKO phenotype in the form of reduced pigmentation, blood accumulation on the yolk, deformed vasculature and a lack of circulation. The Non Tg sibling control (genotype *tdpl +/-;tdp43 -/-*) and *MM tdpl ^RC^ ^ex^*^2^*^:exorh-GFP-PA^* sibling control (*MM tdpl ^RCex^*^2^*^:exorh-GFP-PA^ /-; tdp43 +/-)* demonstrating no phenotype. The DKO phenocopy validates *MM tdpl ^RC ex^*^2^*^:exorh-GFP-PA^* as loss of function.

### Double compound heterozygous Marked Mutants recapitulate the DKO phenotypes

Having validated all Marked Mutant alleles in *tdp43* and *tdpl* as loss-of-function alleles, we next sought to determine whether double compound heterozygous individuals, carrying two independently generated marked mutant alleles for each gene (*tdp43* and *tdpl*), would phenocopy the *tdp43/tdpl* DKO. These individuals would carry four distinct secondary markers, enabling their unambiguous identification at 24hpf), before the onset of overt phenotypes. This strategy would allow us to prospectively identify DKO-equivalent individuals at scale, facilitating subsequent phenotypic analyses. To do so we crossed MM tdp43 *^ex^*^3^*^:he^*^1^.^1^*^-RFP-PA^*; MM tdpl *_ex1:ins-GFP-PA_* with MM tdp43 *^ex^*^3^*^:he^*^1^.^1^*^-GFP-PA^*_;_ *MM tdpl ^RC ex^*^2^*^:exorh-GFP-PA^* and sorted individuals at 24hpf into their respective genotypes. Only individuals with all four markers (double compound heterozygous) phenocopied the traditional *tdp43/tdpl* DKO displaying reduced pigmentation, impaired circulation, progressive deformity and embryonic lethality (figure 8). The remaining sibling controls carrying 3 Marked Mutant alleles or less, all appeared WT (figure 8). This confirms that Marked Mutant alleles are loss of function and double compound heterozygous individuals phenocopy traditional DKO.

**Figure 8.**
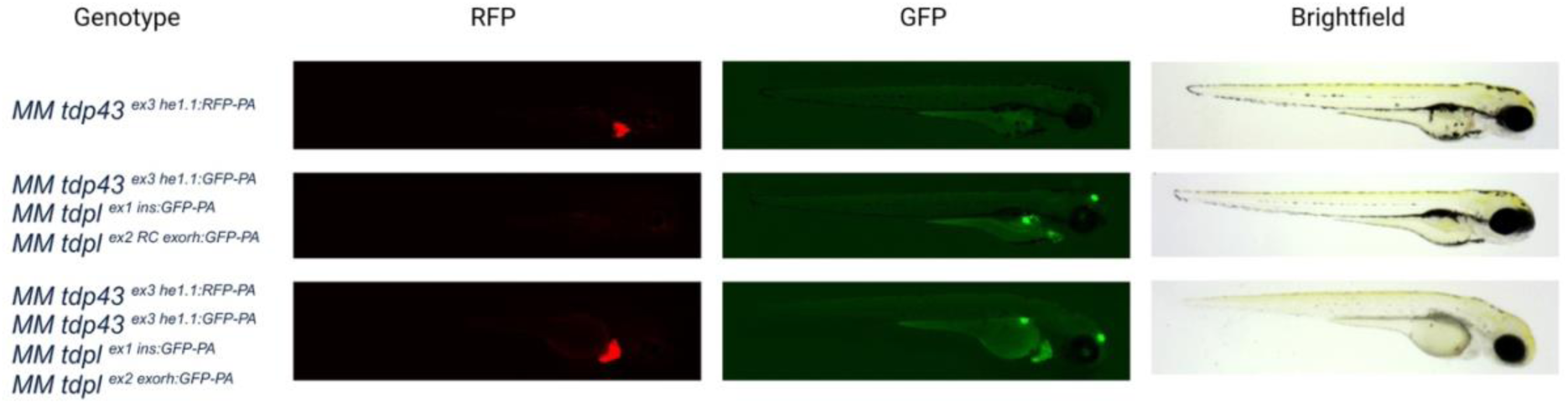
Double compound heterozygous Marked Mutants phenocopy traditional DKO phenotype. Independent double *tdp43/tdpl* marked mutant heterozygous individuals each carrying a different pair of markers (*MM tdp43^ex^*^3^*^:he^*^1^.^1^*^- RFP-PA^; MM tdpl^ex^*^1^*^:ins-GFP-PA^* and *MM tdp43^ex^*^3^*^:he^*^1^.^1^*^-GFP-PA^*_;_ *MM tdpl ^RC ex^*^2^*^:exorh-GFP-PA^*) were crossed together and progeny sorted at 24hpf. F3 progeny carrying a single marker (such as *MM tdp43^ex^*^3^*^:he^*^1^.^1^*^-RFP-PA^*) exhibited no phenotype at 3dpf. Progeny carrying 3 separate markers (such as *tdp43^ex^*^3^*^:he^*^1^.^1^*^-GFP-PA^_/_+*; *MM tdpl ^RC ex^*^2^*^:exorh-GFP-PA^/ tdpl^ex^*^1^*^:ins-GFP-PA^*) exhibited no phenotype. Only individuals carrying all four markers at 3dpf (*MM tdp43^ex^*^3^*^:he^*^1^.^1^*^-RFP-PA^/ tdp43^ex^*^3^*^:he^*^1^.^1^*^-GFP-PA^; MM tdpl ^RC ex^*^2^*^:exorh-GFP-PA^/ tdpl^ex^*^1^*^:ins-GFP-PA^*) developed the classical DKO phenotype in the form of reduced pigmentation, blood accumulation on the yolk, deformed vasculature and a lack of circulation. This demonstrates that double compound heterozygous Marked Mutants are loss of function and phenocopy their respective DKO phenotype.

### Validation of on-target integration is essential for establishing Marked Mutant lines

During characterisation of the *tdpl* lines, we identified two important technical considerations. Firstly, the *MM tdpl^ex^*^1^*^:ins-BFP-PA^* founder generated F1 progeny carrying pancreatic BFP expression both with and without intact insertion junctions. Only animals carrying intact on-target integrations (confirmed by junction PCR analysis) displayed loss-of-function phenotypes, demonstrating that independent on-target and off-target insertion events can coexist within the germline of a single founder. This observation highlights the importance of validating insertion junctions during founder screening ahead of subsequent stable line establishment. Consequently, F1 adults were genotyped following fin biopsy, and only animals carrying intact on-target integrations were retained for propagation. Marked Mutant alleles should then be validated by confirming Mendelian inheritance of the fluorescent marker in the subsequent F2 generation, after which fluorescence can be used for routine genotype identification without molecular genotyping.

### Maternal fluorescence in *tdpl* Marked Mutants is dependent on secondary marker choice

During characterisation of the *tdpl* Marked Mutant lines, we observed maternal fluorescence that varied depending on the secondary marker used. Both insulin promoter-driven *tdpl* Marked Mutant alleles exhibited strong maternal fluorescence, with offspring from female, but not male *MM tdpl ex1:ins GFP/BFP* adults displaying widespread GFP or BFP fluorescence from fertilisation, irrespective of whether they inherited the Marked Mutant allele (Figure 9A–B). This fluorescence gradually dissipated by approximately 2 dpf, suggesting maternal deposition of the fluorophore.

**Figure 9.**
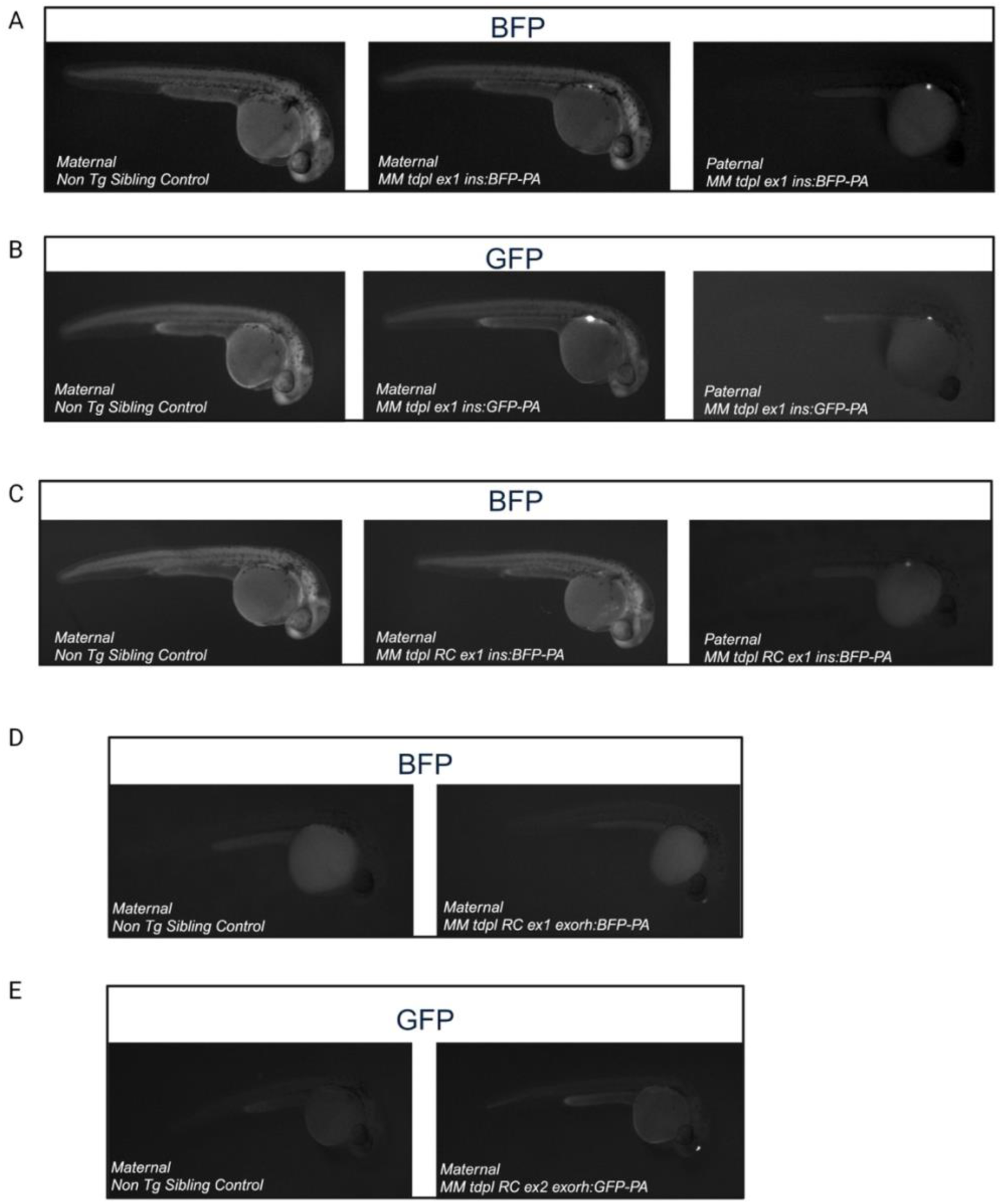
Secondary marker choice influences maternal fluorescence in *tdpl* Marked Mutants. **A-B.** 30hpf *tdpl* marked mutant alleles utilising the insulin promoter to drive BFP (A) or GFP (B), lead to maternal deposition of the fluorophore in their progeny. Maternal *tdpl^ex^*^1^*^:ins-BFP-PA^* and *tdpl^ex^*^1^*^:ins-GFP-PA^* outcrosses show widespread fluorophore deposition in both their MM and non Tg progeny. No fluorophore deposition occurs in paternal outcrosses. **C** 30hpf *tdpl ^RC ex^*^1^*^:ins-BFP-PA^* marked mutants lead to maternal deposition of the fluorophore in their progeny. Maternal *tdpl ^RC ex^*^1^*^:ins-BFP-PA^* outcrosses show widespread fluorophore deposition in both their MM and non Tg progeny despite the secondary marker being inserted into *tdpl* in its reverse orientation. No fluorophore deposition occurs in paternal outcrosses. This demonstrates that secondary marker orientation does not prevent maternal deposition of the fluorophore in *MM tdpl ^RC ex^*^1^*^:ins-BFP-PA^*. **(D–E)** Representative images of 30 hpf progeny from maternal outcrosses of *MM tdpl ^RC ex^*^1^*^:exorh-BFP-PA^* **(D)** and *MM tdpl ^RC ex^*^2^ *^exorh-GFP-PA^* **(E)** heterozygotes. Widespread maternal fluorescence was not detected in progeny from the exon 1 exorh line and was barely detectable in progeny from the exon 2 exorh line under the imaging conditions used. All images were acquired using identical exposure and illumination settings.

We reasoned that maternal fluorescence might arise from interactions between the secondary marker cassette and endogenous *tdpl* regulatory elements, potentially influenced by cassette orientation, promoter choice or proximity to the endogenous promoter. To investigate this, we compared maternal fluorescence across the different *tdpl* Marked Mutant alleles. Reversing the orientation of the insulin-driven BFP cassette in exon 1 (*MM tdpl ^RC ex^*^1^*^:ins-BFP-PA^)* did not prevent maternal fluorescence, with all offspring from female heterozygotes exhibiting widespread BFP expression irrespective of genotype (Figure 9C). This suggests that cassette orientation alone does not determine maternal fluorescence.

In contrast, *tdpl* Marked Mutant alleles carrying pineal gland (exorh)-driven secondary markers (*MM tdpl ^RC^ ^ex^*^1^*^:exorh-BFP-PA^*and *MM tdpl ^RC^ ^ex^*^2^*^:exorh-GFP-PA^*alleles) exhibited substantially reduced maternal fluorescence. Maternal fluorescence was barely detectable in the *tdpl exon 2 exorh allele* and was not detected in the *exon 1 exorh allele* (Figure 9D–E). Importantly, both *exorh* cassettes were inserted in the reverse-complement orientation, as was the insulin-driven BFP cassette, suggesting that promoter choice rather than orientation is a major determinant of maternal fluorescence.

Together, these findings demonstrate that maternal fluorescence in *tdpl* Marked Mutants is strongly influenced by secondary marker selection. Although the precise mechanism remains unclear, interactions between the insulin secondary marker and endogenous *tdpl* regulatory elements may contribute to maternal fluorophore expression. Our findings highlight the importance of secondary marker choice when generating Marked Mutants, particularly for genes with substantial maternal expression, where widespread maternal fluorescence could interfere with early visual genotyping.

### Marked Mutant allele recovery is comparable to traditional CRISPR-Cas9 mutant pipelines

When we pooled the data across all Marked Mutant lines, approximately 30% of injected F0 animals subsequently transmitted validated on-target Marked Mutant alleles through the germline (table 2). Thus, only relatively small numbers of injected adult fish are required for founder screening, representing a workload broadly comparable to that associated with conventional CRISPR–Cas9 zebrafish mutant pipelines^33^. Further analysis of the pooled data from all propagated marked mutant alleles revealed that 4/11 individual founder lines showed some evidence of off target integration of the secondary marker gene within the founder germ line, based on fin biopsy of the F1 individuals. As founders are so common, we would recommend only raising F1 from founders which do not show evidence for off target integration (from F1 embryos). However, as a precaution we recommend confirming secondary marker junctions are present in the F1 adults via traditional fin biopsy, only propagating individuals with on target insertion and monitoring mendelian inheritance in the subsequent generations before switching to full visual genotyping.

**Table 2.** Founder screening recovery rates for each Marked Mutant line. Recovery rates of each marked mutant line, with the initial number of raised injected F0 fish screened and the proportion of those that produced F1 positive for the on-target insertion of the secondary marker. ∼30% of raised injected individuals are founders for on target insertion.

| Gene (Exon) | Secondary Marker | On Target Insertion Founders |
|---|---|---|
| *mitfa (ex4)* | *exorh-gfp* | 3 out of 4 |
| *tdp43 (ex3)* | *he1.1-rfp* | 3 out of 8 |
| *tdp43 (ex3)* | *he1.1-gfp* | 1 out of 10 |
| *tdpl (ex1)* | *ins-bfp* | 1 out of 5 |
| *tdpl (ex1)* | *ins-gfp* | 3 out of 7 |
| *tdpl (ex1)* | *rc ins-bfp* | 1 out of 7 |
| *tdpl (ex1)* | *rc exorh-bfp* | 3 out of 11 |
| *tdpl (ex2)* | *rc exorh-gfp* | 1 out of 4 |

Finally, many secondary markers described in the literature are available in only a limited range of fluorescent colours or are contained within plasmid backbones that can be challenging to propagate, such as Tol2 Destination vectors^15^. Moreover, the suitability of individual secondary markers depends on the experimental context. For example, the markers used in this study may be unsuitable for researchers investigating different organs/tissues to the CNS, where unobstructed visualisation of the pancreas or yolk may be required. To increase the flexibility and accessibility of secondary marker selection, we generated, where not already available, plasmids encoding RFP, GFP, and tagBFP fluorescent reporters for the hatching gland, pancreas, heart, eyes, and pineal gland (Table 1). Importantly, these plasmids can be propagated using conventional competent cells, facilitating their routine use and distribution.

## Discussion

We have described and validated a strategy for generating stable loss-of-function zebrafish mutants that can be genotyped rapidly and non-invasively by simple fluorescence stereomicroscopy. Our Marked Mutant approach enables simultaneous gene disruption and fluorescent labelling of mutant alleles in a single targeted integration event. Once Marked Mutant lines have been established, the need for routine PCR-based genotyping is eliminated. The Marked Mutant workflow is simple and inexpensive, with correctly targeted insertions transmitted through the germline by approximately 30% of fluorescence-selected F0 adults screened. Thus, relatively few selected F0 adults need to be screened to identify germline-transmitting founders, consistent with the modest screening requirements reported for conventional CRISPR–Cas9 zebrafish mutagenesis pipelines^33^. Moreover, all Marked Mutant alleles examined phenocopied their respective conventional loss-of-function mutants, validating exon-targeted insertion of secondary markers as an effective mutagenesis strategy.

Because biallelic mutants can be identified unambiguously and rapidly by visual inspection before overt phenotypes emerge, embryonic lethal mutants can be incorporated into scalable chemical screening paradigms^34^. Rapid discrimination of wild-type, heterozygous and biallelic mutant individuals also simplify experimental design and colony management. The ability to combine multiple orthogonal secondary markers further facilitates the generation and maintenance of double and triple mutant combinations, an important advantage in zebrafish where many genes exist as duplicated paralogues^7^. Moreover, if the Marked Mutant strategy is combined with robotic embryo sorters, this has the potential to fully automate the genotyping process in zebrafish^35^.

An additional advantage of our approach is its impact on animal husbandry and welfare. Because genotyping of Marked Mutant alleles can be performed as early as 24hpf, only animals of the desired genotype need to be raised to adulthood, substantially reducing tank usage and associated maintenance costs. Non-invasive visual genotyping at pre 5dpf stages also minimises the number of genetically modified animals progressing to protected stages and removes the requirement for invasive adult fin clipping, a regulated procedure in many jurisdictions^2^. Given that raising excess animals and subsequent fin biopsy constitute a substantial proportion of regulated procedures in animal research, widespread implementation of visually genotyped zebrafish alleles could significantly reduce the overall procedural burden associated with animal experimentation for biomedical research^2^.

Related genome editing approaches enabling non-invasive visual genotyping in zebrafish have previously been reported including the UFlip, CRIMP and TICIT methods^19–21^. These methods typically rely on intronic targeting strategies (UFlip/CRIMP) in which fluorescent reporters are linked to the target genes promoter and downstream transcription is terminated through the addition of transcriptional stop elements^19,21^. Although highly effective at inhibiting target gene function, with UFlip also enabling cell-type-specific knockout, such strategies generally require cloning (UFlip) and depend on efficient intron targeting (UFlip/CRIMP)^19,21^. The latter can be challenging because intronic sequences are often poorly conserved and enriched for repetitive or AT-rich regions that complicate CRISPR–Cas9 targeting. By contrast, our approach is entirely PCR-based and targets coding exons, which are typically more conserved and more readily amenable to genome editing. Importantly, in our strategy, fluorophore expression is driven independently of the endogenous transcriptional pattern of the target gene and is instead restricted to discrete anatomical structures. This may be advantageous for live imaging applications, where broader fluorophore expression, as can occur with UFlip and CRIMP, could complicate visualization. Furthermore, when fluorophore expression is linked to the target gene promoter, genotype identification may be more challenging for genes with low or conditional expression levels, and maternal fluorophore contribution may also occur. By contrast, our independently driven secondary markers allow genotype identification irrespective of target gene expression levels, while potential maternal fluorescence associated with certain targets may be minimized through the selection of alternative markers.

TICIT is perhaps the most comparable strategy to our approach, although the two differ in their integration workflows. TICIT involves two sequential integration steps: the initial insertion of a phiC31 landing site into the target gene using CRISPR–Cas9, followed by site-specific integration of a fluorescent reporter cassette using phiC31 integrase^20^. In contrast, the Marked Mutant approach requires only a single targeted integration step. Furthermore, approximately 30% of screened F0 animals transmitted on-target Marked Mutant alleles, compared with founder frequencies of 6–33% for the initial TICIT landing-site integration^20^.

A limitation we encountered with our approach was the occurrence of founders producing F1 progeny with both off and on target insertions of the secondary marker. Only the latter are loss of function. From all propagated alleles we found 4/11 founders had evidence of F1 with and without on target insertions. The issue of off target integration of a secondary marker is not a unique issue to our approach as it has been reported in approaches to generate zebrafish floxed alleles for cell type specific KO^36^. Of note we have also encountered this phenomena when generating own cell type specific KO lines using the UFlip approach^37^. A simple solution is to propagate only founders that show no evidence of off-target insertions in their F1 offspring, as correctly targeted Marked Mutant founders are recovered at a sufficiently high frequency. Nevertheless, we recommend confirming correct junction integration in F1 adults and verifying Mendelian inheritance of the secondary marker in the F2 generation.

Beyond these technical considerations, our approach offers the potential to bring the complex combinatorial genetics routinely performed in Drosophila to a vertebrate model system. To facilitate this, we have assembled a library of spectrally distinct secondary markers that enable visual genotype identification from 24hpf (table 1). Future work could expand this library to include markers detectable before 24hpf, enabling the study of mutant phenotypes arising during the earliest stages of development.

In summary, we describe and demonstrate a rapid and scalable strategy for generating visually identifiable loss-of-function zebrafish alleles, which we refer to as Marked Mutants. Our method retains the simplicity and efficiency of conventional CRISPR-based mutagenesis pipelines while enabling non-invasive genotyping by direct visual inspection, reducing experimental burden and enhancing animal welfare in zebrafish reverse genetics. Furthermore, our approach facilitates combinatorial genetics by enabling the non-invasive identification of double homozygous mutants, before overt phenotypes emerge.

## Materials and Methods

### Zebrafish Husbandry

Adult zebrafish (*Danio rerio*) were housed and maintained in accordance with standard procedures in the Queen’s Medical Research Institute zebrafish facility, University of Edinburgh. All experiments were performed in compliance with the UK Home Office, according to its regulations under project licences 70/8436, PP5258250 and PP3290955. Adult zebrafish were subject to a 14/10 h light/dark cycle. Embryos were produced by pairwise matings and were staged according to days post-fertilisation (dpf). Embryos were raised at 28.5°C, with 50 embryos (or less) per 10 cm petri dish in ∼45 ml of 10 mM HEPES-buffered E3 embryo medium or conditioned aquarium water with methylene blue.

### Plasmids

The GFP pineal gland marker *exorh:GFP-PA* was purchased from Addgene (plasmid number #213514)^38^. The BFP pineal gland marker *exorh:BFP-PA* was generated as a Gblock (IDT) and converted into a middle entry clone using BP clonase II with plasmid pDONR221 (Invitrogen) and gateway technology^15^ using primers F’ GGGGACAAGTTTGTACAAAAAAGCAGGCTGCTCAGCTGGCAGTACTACC and R 5’ GGGGACCACTTTGTACAAGAAAGCTGGGTTCACAAAAGCCATTTTTATTG. The RFP pineal gland marker *exorh:RFP-PA* was generated via fast cloning (see table 3) GFP hatching gland plasmid, *he1.1:GFP-PA* and was generated as a Gblock and converted into a middle entry clone using primers F 5’ GGGGACAAGTTTGTACAAAAAAGCAGGCTTCAACCACTCCAGGCATAGCT and R 5’ GGGGACCACTTTGTACAAGAAAGCTGGGTAAAAAACCTCCCACACCTCC.

**Table 3.**
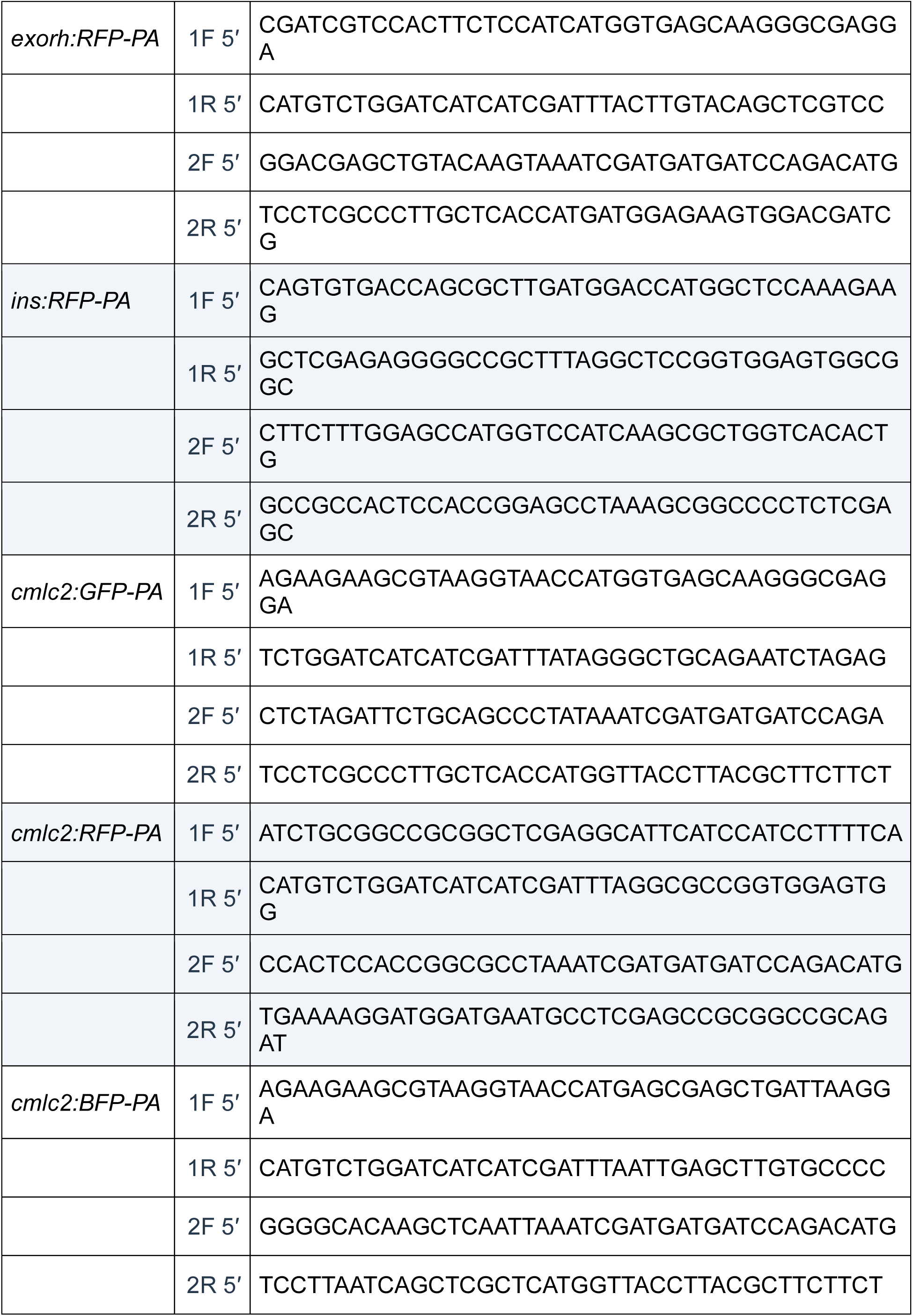
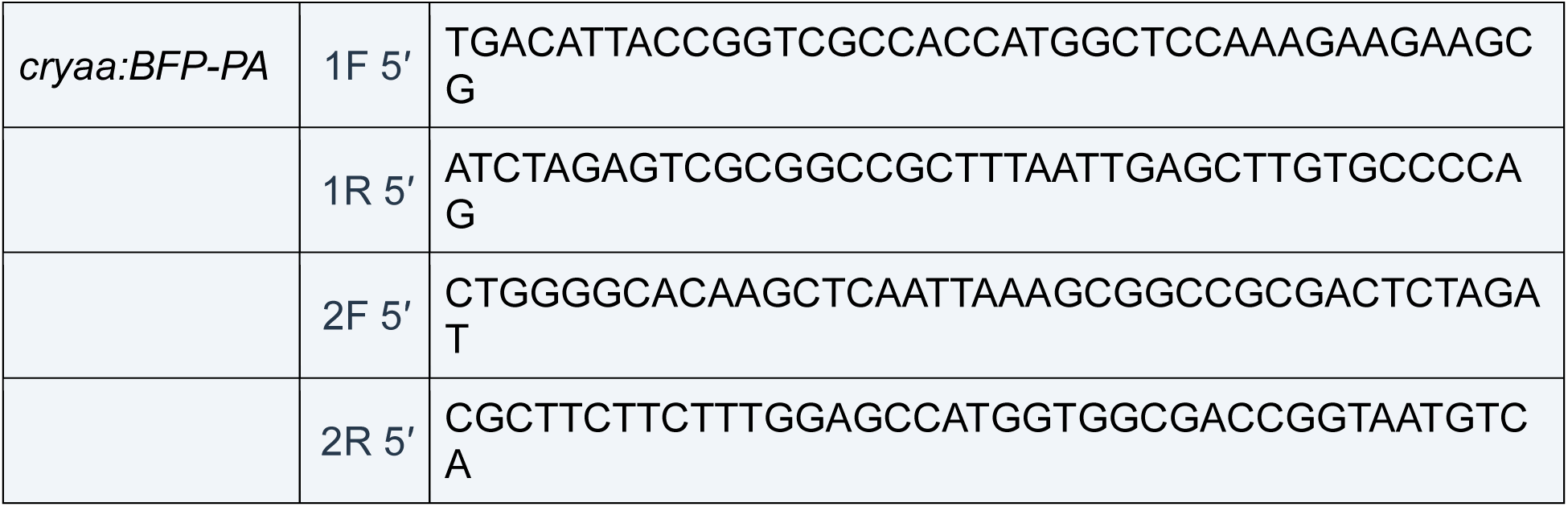
Fast cloning primer sequences.

The RFP hatching gland marker *he1.1 RFP-PA* was generated in the same fashion but with primers F 5’ GGGGACAAGTTTGTACAAAAAAGCAGGCTTCAACCACTCCAGGCATAG and R 5’ GGGGACCACTTTGTACAAGAAAGCTGGGTAAAAAACCTCCCACACCTCC. The *He1.1:BFP-PA* was amplified from a UFLIP plasmid *pUFlip-floxed2A-mRFP-; HE1A:BFP -0*^19^ purchased from Addgene (plasmid number #180007) and converted into a middle entry clone using primers F 5’ GGGGACAAGTTTGTACAAAAAAGCAGGCTTCAACCACTCCAGGCATAGC and R 5’ GGGGACCACTTTGTACAAGAAAGCTGGGTTCACAAAAGCCATTTTTATT.

The GFP pancreas marker, *ins:GFP-PA* was a kind gift from Angeleen Fleming (University of Cambridge). A ∼1kb fragment of the promoter was cloned from genomic zebrafish DNA to include the promoter region that had previously been used to generate published *ins:EGFP* lines^31^. The PCR fragment was ligated into the TOPO TA cloning vector (Thermo Fisher Scientific) then subcloned into a destination vector using restriction enzymes. The *ins:GFP-PA* destination vector was converted into a middle entry clone using primers F 5’ GGGGACAAGTTTGTACAAAAAAGCAGGCTAGATCTTTTCGCTAATCAGCC and R 5’ ACCACTTTGTACAAGAAAGCTGGGTCTTGTTTATTGCAGCTTATA. The BFP pancreas marker, *ins:BFP-PA* was generated as a Gblock and converted into a middle entry clone using primers F 5’ GGGGACAAGTTTGTACAAAAAAGCAGGCTTTTCGCTAATCAGCCCACAGT and R 5’ GGGGACCACTTTGTACAAGAAAGCTGGGTAAAAAACCTCCCACACCT. The RFP pancreas marker was generated by fast cloning (see table 3). Heart marker plasmids were all made via fast cloning (see table 3). GFP/mCherry lens marker plasmids were made by converting destination vectors into middle entry clones using primers F 5’ GGGGACAAGTTTGTACAAAAAAGCAGGCTAAAATTGTGAAATGCACAGA and R 5’ GGGGACCACTTTGTACAAGAAAGCTGGGTTAAGATACATTGATGAGTTT (for *cryaa:GFP-PA*) and F 5’ GGGGACAAGTTTGTACAAAAAAGCAGGCTAAAATTGTGAAATGCACAGAA and R 5’ GGGGACCACTTTGTACAAGAAAGCTGGGTTAAGATACATTGATGAGTTTG (for cryaa:mcherry-PA).

### Fast Cloning

FastCloning was performed as described by Li et al^39^.The target DNA insert and plasmid vector were amplified separately using Q5 (NEB), with primers designed to introduce 20bp overlapping sequences. The PCR products were mixed and treated with DpnI (NEB) at 37°C for 1 hour to remove parental plasmid DNA before direct transformation into competent E. coli Top10cells (Invitrogen). Positive clones were confirmed by whole plasmid sequencing. Primers for each fast clone are located in table 3.

### Crispr KI design and donor DNA generation

CrRNAs were designed to target restriction enzyme sites whereby the Cas9 cut site crossed the restriction enzyme recognition site meaning restriction digest could be used to assay mutagenesis as previously described^40^. Early exons, especially those coding for domains were targeted. 2nMol of each CrRNA and 5nMol of TracrRNA without modifications (Sigma/Merck), were resuspended in 100μl and 250μl of nuclease free water respectively and frozen at -20°C until the day of injection. CrRNA sequences are outlined in table 4. Knock-In was performed using the PCR based Gene Tagging method as previously described^16^, briefly, primers were designed containing 40bp homology arms to the target insertion location with 20bp sequence used to amplify the secondary marker plasmid DNA. The homology arms were designed to insert the secondary marker directly at the Cas9 double strand cut site. Primers were ordered through IDT, all were biotinylated at the 5’ end and had phosphorothioate bonds in the first 5 nucleotides in the 5ʹ-end. Donor DNA primers were used to amplify the entire secondary marker (promoter, fluorophore with ATG start/TAA stop sequences and polyA tail) with target gene homology arms, using X2 Q5 proof reading polymerase (NEB). The PCR product was run on a 1% agarose gel to confirm the correct band size, purified by DNA gel extraction (Monarch/NEB), eluted in 6μl nuclease free water and frozen at -20°C until the day of injection. Filter tips were used for all steps. All donor DNA primers can be found in table 5.

**Table 4.**
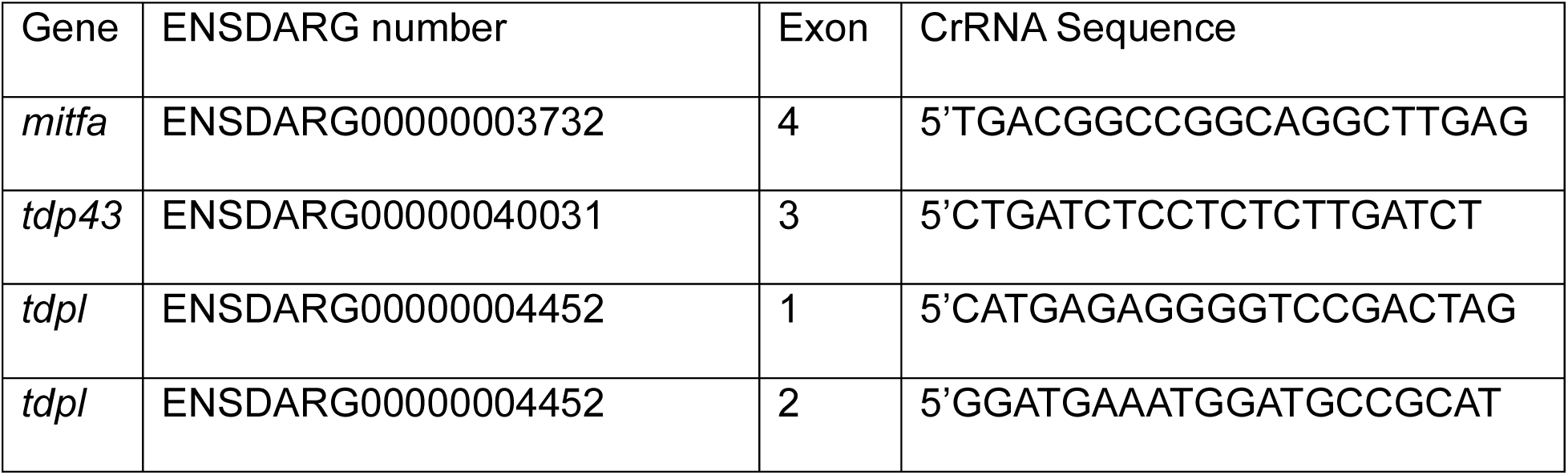
CrRNA sequences.

**Table 5.**
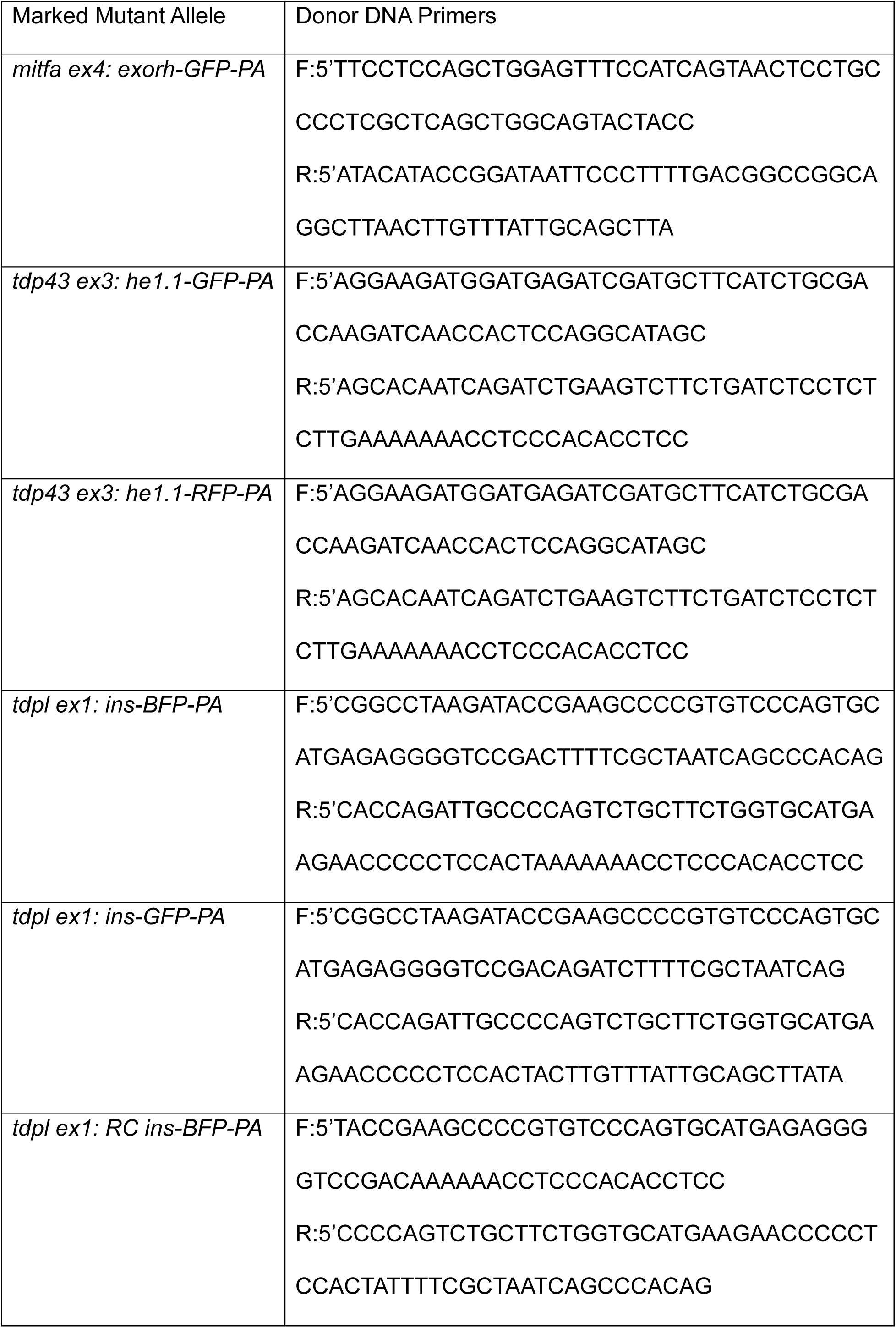
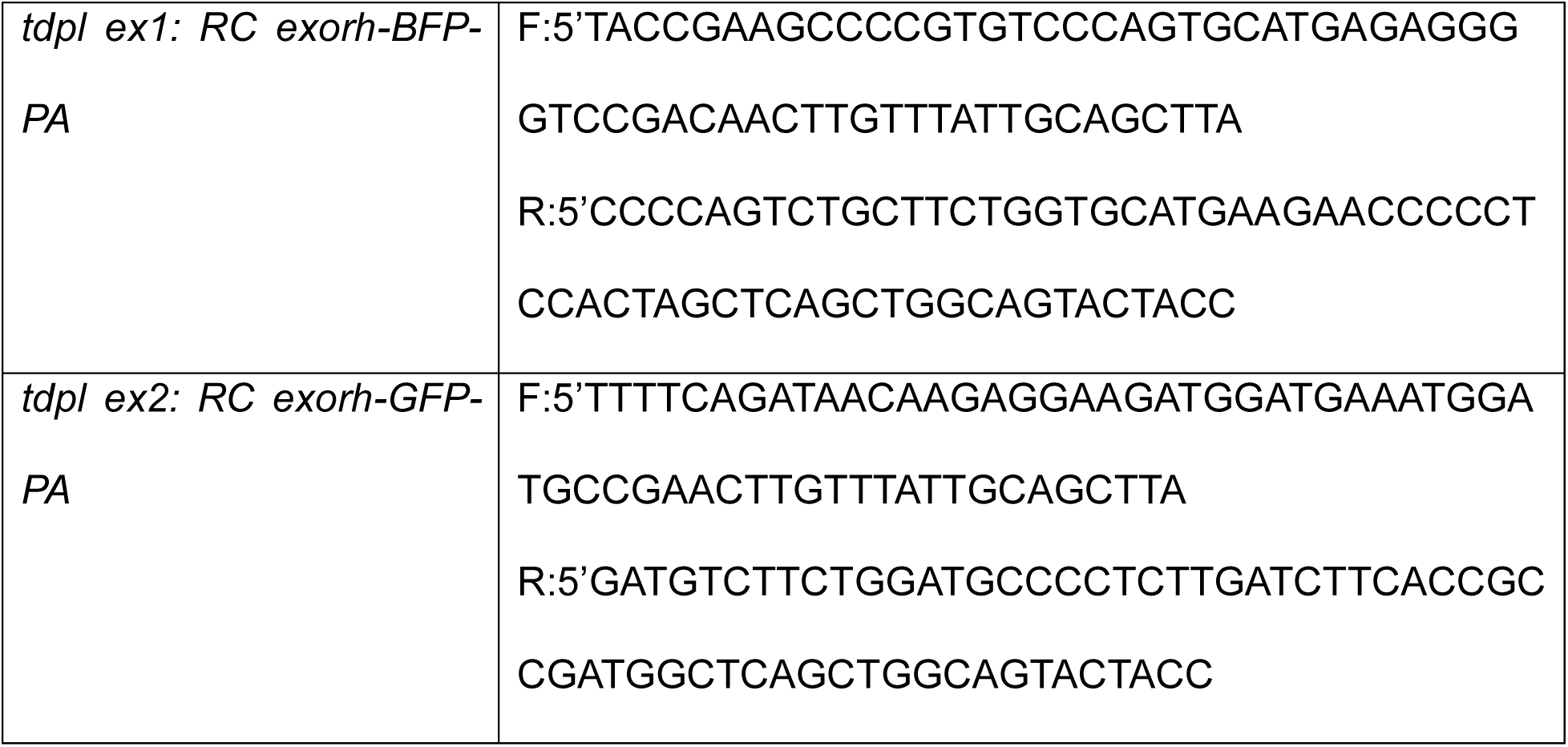
All primers used to generate donor DNA for knock in.

### Crispr/Cas9 injections

On the day of injection, 2 µl of crRNA (20 µM) was combined with 2 µl of tracrRNA (20 µM) in a PCR tube by gentle pipetting. The RNA mixture was annealed by heating to 95 °C for 5 minutes, followed by cooling on ice^40,41^. Subsequently, 200ng of purified PCR product donor DNA was added together with 1µl Cas9 protein (M0646M, NEB), 1µl phenol red, and 0.2 µl of 7.4 µM NU7441 (APExBio), before the solution was brought to a final volume of 8 µl using nuclease-free water. The mixture was gently pipetted to ensure thorough mixing. The injection mix was then incubated at 37 °C for 5 minutes, as previously described^42^, and subsequently cooled on ice. Filter tips were used throughout all procedures. Wild-type embryos were injected with 1nl of the mixture directly into the cell at the one-cell stage. Only injected individuals with widespread expression of fluorophore within the secondary marker organ, were analysed by PCR and raised for founder screening.

### Restriction Digest

At 24hpf, CrRNAs were tested for mutagenic activity via restriction digest^40^. Whole embryos were added to individual wells of a PCR plate and homogenised by incubating in 100µl 50mM NaOH for ten minutes at 95°C then vortexed briefly. The NaOH was neutralised with 10µl 1M Tris-HCL pH8.0 and vortexed again to mix. A region of genomic material ∼150-500bp in size was amplified from the embryo homogenate using BiomixRed (Bioline). Post PCR, 1μl of the respective restriction enzyme (table 6) was added to the PCR product and incubated following the manufacturer’s instructions (NEB) and then run on a 2% agarose gel. Restriction digest was conducted on 4 uninjected and 4 injected individuals. DNA samples from different embryos were not pooled.

**Table 6.**
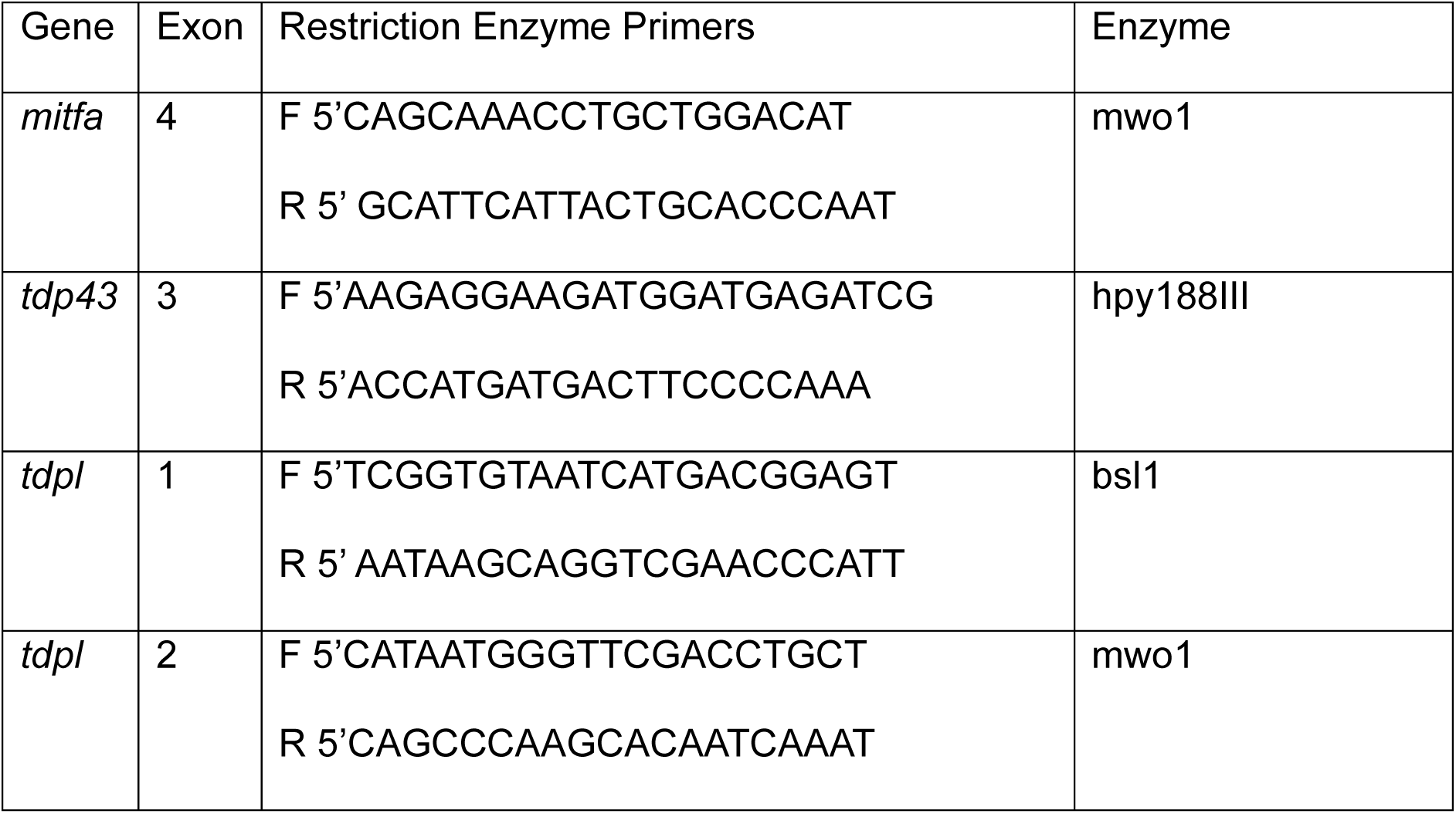
Restriction enzyme primers.

### Founder screening

Injected individuals (F0) with widespread expression of the secondary marker were raised to 3 months of age. Founders were identified by crossing individual injected fish (F0) to WT adults (pair mating) and then inspection of F1 progeny for the secondary marker. As alleles could potentially be rare, only clutches with >50 fertile F1 embryos were included in the subsequent analysis. F1 were inspected at 24hpf and 48hpf for development of the secondary marker. F1 positive for the transgene were analysed by junction PCR analysis to confirm successful on target integration using primers ideally binding to exonic material where possible, amplifying a region of 400-800bp. Due to potential indels, primers were designed to bind at least 100bp away from the integration site. Where possible, 4 non transgenic and 4 transgenic embryos were analysed using a pair of primers (outlined in table 7) for presence or absence of a PCR signal. PCR bands denoting on target integration were sequenced by traditional Sanger sequencing.

**Table 7.**
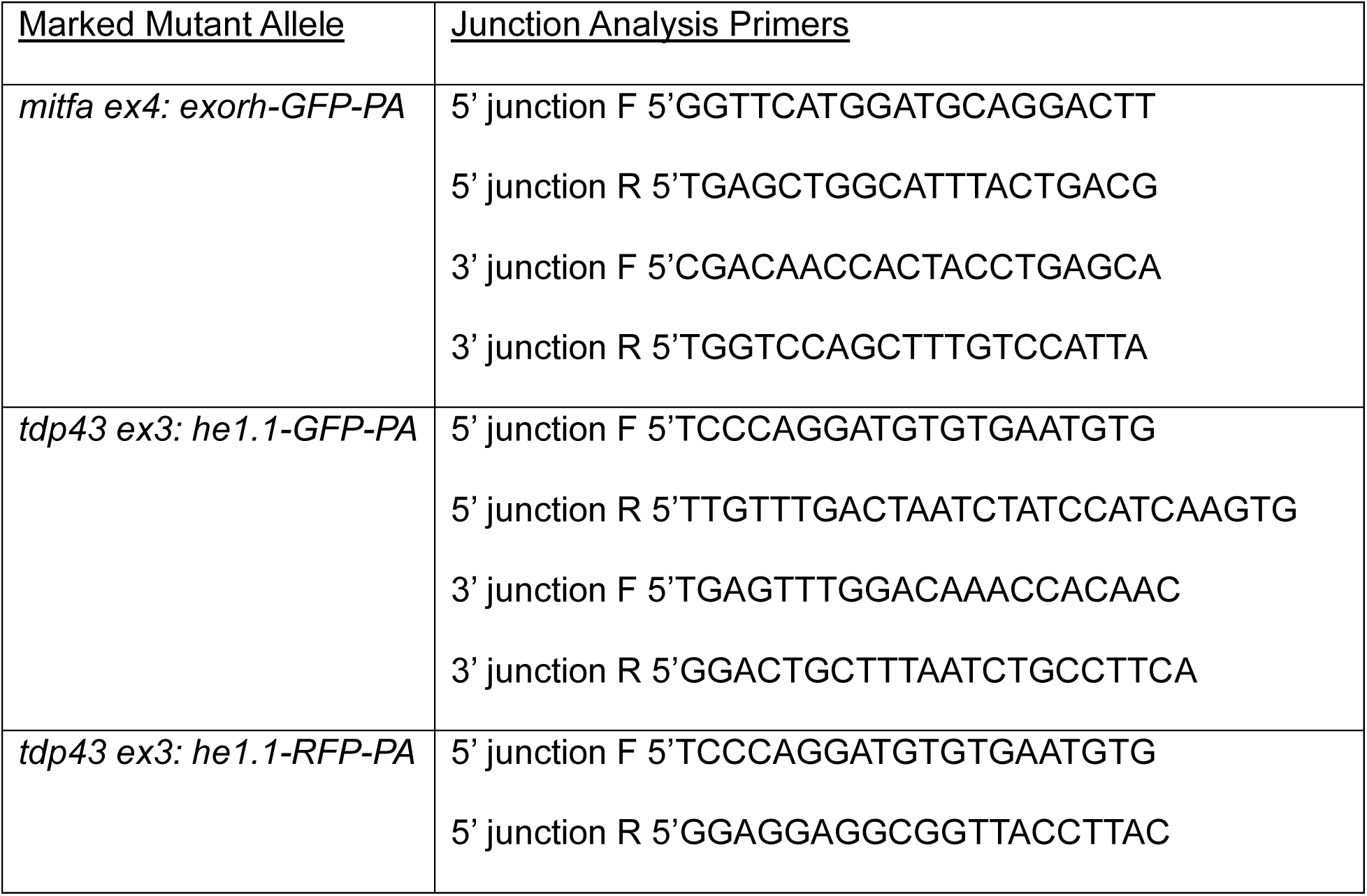
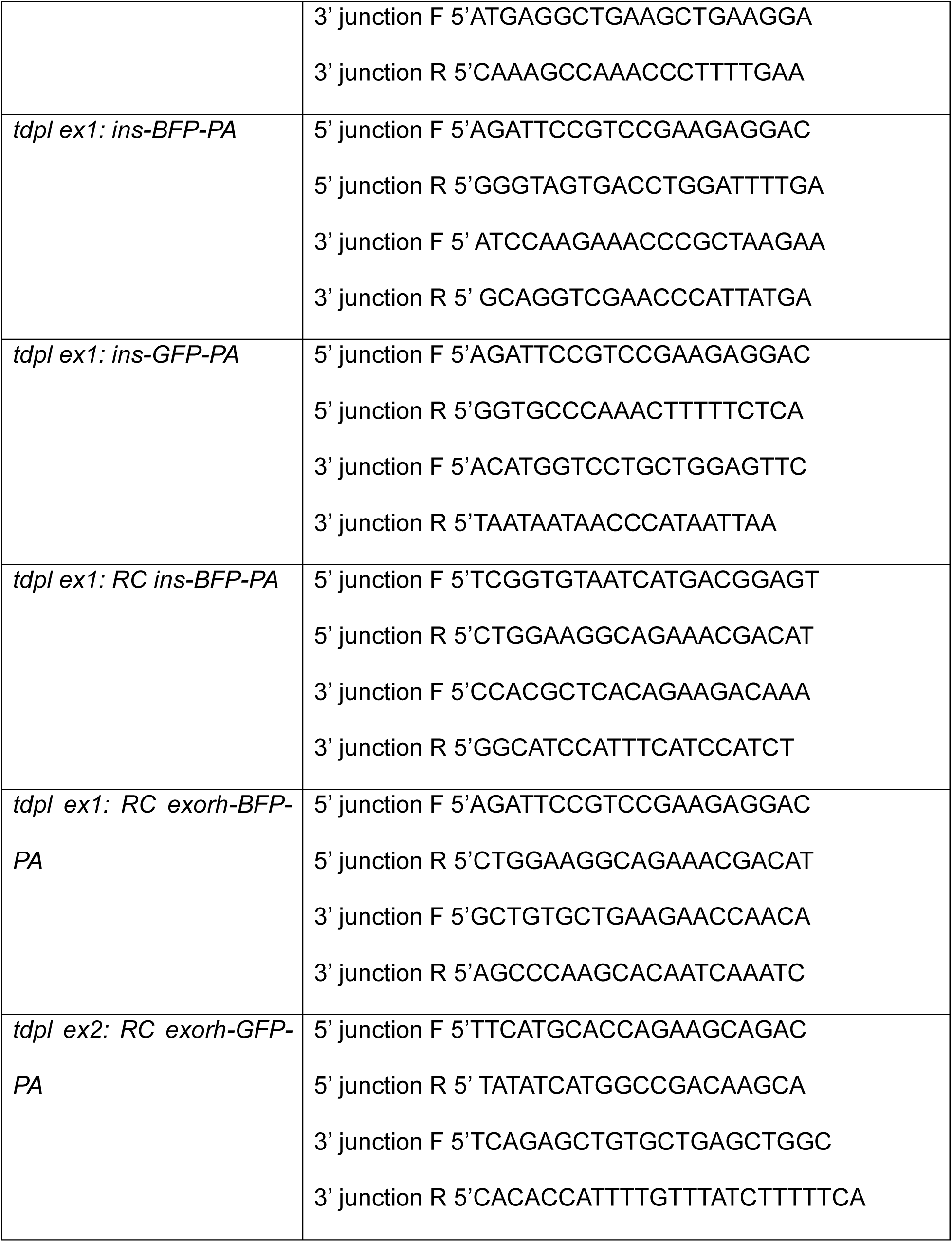
List of junction PCR primers used for junction analysis.

### Conventional Crispr/Cas9 stable mutants used in the study

Loss of function mutants for TDP43 (10bp del exon 3) and TDPL (56bp del +18bp ins exon 1) were generated by CRISPR-Cas9-based gene editing, using CrRNAs targeting sequence 5’ TGGGGAAGTCATCATGGTGC (TDP43) and 5’ CATGAGAGGGGTCCGACTAG (TDPL) respectively^43^. Each embryo was genotyped post hoc after data analysis using TDP43 primers F 5’ GCGACCAAGATCAAGAGAGG and 5’ R ACCACGGGTGTATGGGTTAT followed by an msl1 digest. TDPL was genotyped using primers 5’ F TCGGTGTAATCATGACGGAGT and 5’ AATAAGCAGGTCGAACCCATT and genotyped by band shift on a gel due to the large size of the indel.

## Notes

### Competing Interest Statement

The authors have declared no competing interest.

## References

1. Xing, L. et al. Rapid and Efficient Zebrafish Genotyping Using PCR with High-resolution Melt Analysis. J. Vis. Exp. e51138 (2014) doi:10.3791/51138.

2. Sneddon, L. U. et al. Pain management in zebrafish. Lab. Anim. 58, 261–276 (2024).

3. Wilkinson, R. N., Elworthy, S., Ingham, P. W. & van Eeden, F. J. M. A method for high-throughput PCR-based genotyping of larval zebrafish tail biopsies. Biotechniques 10.2144/000114116 (2013) doi:10.2144/000114116.

4. Lambert, C. J. et al. An automated system for rapid cellular extraction from live zebrafish embryos and larvae: Development and application to genotyping. PLoS One 13, e0193180 (2018).

5. Venditti, M. et al. A minimally invasive fin scratching protocol for fast genotyping and early selection of zebrafish embryos. Scientific Reports 2022 12:1 12, 22597- (2022).

6. Rasouli, S. J. et al. The flow responsive transcription factor Klf2 is required for myocardial wall integrity by modulating Fgf signaling. Elife 7, (2018).

7. Heppert, J. K. et al. Conserved roles for Hnf4 family transcription factors in zebrafish development and intestinal function. Genetics 222, (2022).

8. Bose, P., Armstrong, G. A. B. & Drapeau, P. Neuromuscular junction abnormalities in a zebrafish loss-of-function model of TDP-43. J. Neurophysiol. 121, 285–297 (2019).

9. Schmid, B. et al. Loss of ALS-associated TDP-43 in zebrafish causes muscle degeneration, vascular dysfunction, and reduced motor neuron axon outgrowth. Proc. Natl. Acad. Sci. U. S. A. 10.1073/pnas.1218311110 (2013) doi:10.1073/pnas.1218311110.

10. Larbalestier, H. et al. GCH1 Deficiency Activates Brain Innate Immune Response and Impairs Tyrosine Hydroxylase Homeostasis. Journal of Neuroscience 42, 702–716 (2022).

11. Keatinge, M. et al. Unexpected phenotypic and molecular changes of combined glucocerebrosidase and acid sphingomyelinase deficiency. 10.1242/dmm.049954 (2023) doi:10.1242/dmm.049954.

12. Keatinge, M. et al. Glucocerebrosidase 1 deficient Danio rerio mirror key pathological aspects of human Gaucher disease and provide evidence of early microglial activation preceding alpha-synuclein-independent neuronal cell death. Hum. Mol. Genet. 24, 6640–6652 (2015).

13. Millerid, D. E., Cook, K. R. & Scott Hawleyid, R. The joy of balancers. 10.1371/journal.pgen.1008421 (2019) doi:10.1371/journal.pgen.1008421.

14. Xie, X. et al. Silencer-delimited transgenesis: NRSE/RE1 sequences promote neural-specific transgene expression in a NRSF/REST-dependent manner. BMC Biol. 10, (2012).

15. Kwan, K. M. et al. The Tol2kit: A multisite gateway-based construction Kit for Tol2 transposon transgenesis constructs. Developmental Dynamics 236, 3088–3099 (2007).

16. Zhang, Y., Marshall-Phelps, K. & de Almeida, R. G. Fast, precise and cloning-free knock-in of reporter sequences in vivo with high efficiency. Development 150, (2023).

17. Mi, J. & Andersson, O. Efficient knock-in method enabling lineage tracing in zebrafish. Life Sci. Alliance 6, (2023).

18. Levic, D. S., Yamaguchi, N., Wang, S., Knaut, H. & Bagnat, M. Knock-in tagging in zebrafish facilitated by insertion into non-coding regions. Development (Cambridge) 148, (2021).

19. Liu, F. et al. Cre/lox regulated conditional rescue and inactivation with zebrafish UFlip alleles generated by CRISPR-Cas9 targeted integration. Elife 11, (2022).

20. Ma, J. et al. Instantaneous visual genotyping and facile site-specific transgenesis via CRISPR-Cas9 and phiC31 integrase. Biol. Open 13, (2024).

21. Miles, L. B. et al. CRIMP: a CRISPR/Cas9 insertional mutagenesis protocol and toolkit. Nature Communications 2024 15:1 15, 5011- (2024).

22. Lister, J. A., Robertson, C. P., Lepage, T., Johnson, S. L. & Raible, D. W. nacre encodes a zebrafish microphthalmia-related protein that regulates neural-crest-derived pigment cell fate. Development 126, 3757–3767 (1999).

23. Kemmler, C. L. et al. Next-generation plasmids for transgenesis in zebrafish and beyond. Development (Cambridge) 150, (2023).

24. Mosimann, C. et al. Site-Directed Zebrafish Transgenesis Into Single Landing Sites With the phiC31 Integrase System. 10.1002/dvdy.23989 (2013) doi:10.1002/dvdy.23989.

25. Solchenberger, B., Russell, C., Kremmer, E., Haass, C. & Schmid, B. Granulin knock out zebrafish lack frontotemporal lobar degeneration and neuronal ceroid lipofuscinosis pathology. PLoS One 10.1371/journal.pone.0118956 (2015) doi:10.1371/journal.pone.0118956.

26. Gurberg, T. J. N. et al. Impaired Motor Function in a Zebrafish Stathmin-2 Knockout Model. Dev. Neurobiol. 86, e70054 (2026).

27. Petel Légaré, V., et al. Loss of mitochondrial Chchd10 or Chchd2 in zebrafish leads to an ALS-like phenotype and Complex I deficiency independent of the mitochondrial integrated stress response. Dev. Neurobiol. 83, 54–69 (2023).

28. Braaker, P. N. et al. Activity-driven myelin sheath growth is mediated by mGluR5. Nature Neuroscience 2025 28:6 28, 1213–1225 (2025).

29. Prater, K. E., Latimer, C. S. & Jayadev, S. Glial TDP-43 and TDP-43 induced glial pathology, focus on neurodegenerative proteinopathy syndromes. 10.1002/glia.24096 (2021) doi:10.1002/glia.24096.

30. Asakawa, K., Handa, H. & Kawakami, K. Optogenetic modulation of TDP-43 oligomerization accelerates ALS-related pathologies in the spinal motor neurons. Nat. Commun. 11, 1–16 (2020).

31. Huang, H., Vogel, S. S., Liu, N., Melton, D. A. & Lin, S. Analysis of pancreatic development in living transgenic zebrafish embryos. Mol. Cell. Endocrinol. 177, 117– 124 (2001).

32. Goishi, K. et al. αA-crystallin expression prevents γ-crystallin insolubility and cataract formation in the zebrafish cloche mutant lens. Development 133, 2585–2593 (2006).

33. Varshney, G. K. et al. High-throughput gene targeting and phenotyping in zebrafish using CRISPR/Cas9. Genome Res. 25, 1030–1042 (2015).

34. Early, J. J. et al. An automated high-resolution in vivo screen in Zebrafish to identify chemical regulators of myelination. Elife 10.7554/eLife.35136 (2018) doi:10.7554/eLife.35136.

35. Letamendia, A., et al. Development and Validation of an Automated High-Throughput System for Zebrafish In Vivo Screenings. PLoS One 7, e36690 (2012).

36. Shin, M. et al. Generation and application of endogenously floxed alleles for cell-specific knockout in zebrafish. Dev. Cell 58, 2614–2626.e7 (2023).

37. Keatinge, M. et al. Mislocalisation of TDP-43 to the cytoplasm of either neurons or oligodendrocytes causes axonopathy and dysmyelination. *bioRxiv* 2025.11.17.688801 (2025) doi:10.1101/2025.11.17.688801.

38. Kemmler, C. L. et al. Next-generation plasmids for transgenesis in zebrafish and beyond. Development (Cambridge) 150, (2023).

39. Li, C. et al. FastCloning: a highly simplified, purification-free, sequence- and ligation-independent PCR cloning method. BMC Biotechnology 2011 11:1 11, 92- (2011).

40. Keatinge, M. et al. CRISPR gRNA phenotypic screening in zebrafish reveals pro-regenerative genes in spinal cord injury. PLoS Genet. 17, e1009515 (2021).

41. Kroll, F. et al. A simple and effective F0 knockout method for rapid screening of behaviour and other complex phenotypes. *bioRxiv* 10.1101/2020.06.04.133462 (2020) doi:10.1101/2020.06.04.133462.

42. Burger, A. et al. Maximizing mutagenesis with solubilized CRISPR-Cas9 ribonucleoprotein complexes. 10.1242/dev.134809 (2016) doi:10.1242/dev.134809.

43. Keatinge, M. et al. Mislocalisation of TDP-43 to the cytoplasm of either neurons or oligodendrocytes causes axonopathy and dysmyelination. *bioRxiv* 2025.11.17.688801 (2025) doi:10.1101/2025.11.17.688801.

